# Tumor architecture shapes polarized malignant states that predict outcome and treatment response in high-grade serous ovarian cancer

**DOI:** 10.64898/2026.05.27.727977

**Authors:** Sarah Nersesian, John Abou-Hamad, Emma M. Durocher, Gaëlle Akiki, Celine Domecq, Athena Southworth, Katrina N. Verey, Hugh H. Deng, Yudith Ramos-Valdes, Liliane Meunier, Manon de Ladurantaye, Trevor G. Shepherd, Diane Provencher, Anne-Marie Mes-Masson, Basile Tessier Cloutier, David P. Cook

**Affiliations:** Cancer Research Program, Ottawa Hospital Research Institute, Ottawa, ON, Canada; Department of Cellular and Molecular Medicine, University of Ottawa, Ottawa, ON, Canada; Department of Pathology, McGill University, Montreal, QC, Canada; Cancer Research Program, Research Institute of the McGill University Health Centre, Montreal, QC, Canada; Centre de recherche du Centre hospitalier de l’Université de Montréal and Institut du cancer de Montréal, Montreal, QC, Canada; Department of Medicine, Université de Montréal, Montreal, QC, Canada; Department of Anatomy and Cell Biology, Western University, London, ON, Canada; The Mary & John Knight Translational Ovarian Cancer Research Program, Verspeeten Family Cancer Centre, London, ON, Canada; Department of Oncology, Schulich School of Medicine & Dentistry, Western University, London, ON; Department of Obstetrics & Gynaecology, Schulich School of Medicine & Dentistry, Western University, London, ON, Canada

## Abstract

High-grade serous ovarian carcinoma (HGSC) is among the most genomically unstable cancers, yet how this diversity relates to malignant-cell states and how they influence treatment response remain unclear. Integrating 1.98 million single-cell transcriptomes across 13 studies, we identify a dominant axis of polarization in malignant secretory cells, from proliferative, progenitor-like (SecA) to quiescent, injury-like (SecB) states that recur across patients and coexist within genetic clones. Spatial profiling of eight whole tumors and tissue microarrays from 378 patients links this polarization to tumor architecture: SecB cells occupy avascular, hypoxic niches, where they adopt HIF/NF-κB survival programs, exclude lymphocytes, and co-organize with macrophages exhibiting similar glycolytic states. Their localization and enrichment in ascites link SecB to metastasis. SecB predicts poor survival across independent cohorts, but preferential benefit from bevacizumab in the ICON7 trial. Together, these findings reveal shared malignant phenotypes across genomically diverse tumors and highlight their potential to guide treatment selection.

## INTRODUCTION

High-grade serous ovarian carcinoma (HGSC) is the most common subtype of ovarian cancer and remains the most lethal gynecological malignancy. Clinical management relies on surgical debulking and combination (platinum/taxane) chemotherapy, with maintenance PARP inhibitors for patients with homologous recombination deficiency (HRD) and frequent use of anti-VEGF antibody bevacizumab^1^. Despite high initial response rates, recurrence of therapy-resistant disease is nearly universal^2^. Bulk transcriptomic subtypes (C1/mesenchymal, C2/immunoreactive, C4/differentiated, C5/proliferative) were defined to stratify patients into biologically and therapeutically distinct groups^3,4^. However, these subtypes have shown inconsistent survival associations across cohorts and often reflect sample composition rather than underlying biology^5–7^. They have also not yet translated into tools for predicting treatment response. No molecular classifier reliably identifies patients who will relapse early after chemotherapy, and while bevacizumab prolongs progression-free survival (PFS) by only a few months in a subset of patients, it rarely improves overall survival (OS)^8^. Beyond HRD, treatment decisions in HGSC remain largely unguided by tumor biology.

Single-cell RNA sequencing (scRNA-seq) has resolved the HGSC tumor microenvironment (TME) in greater detail, characterizing immune phenotypes^9^, stromal subtypes and their prognostic contributions^10^, site-specific immune evasion driven by mutational processes^11^, treatment-associated remodeling under chemotherapy and immunotherapy^12–14^, and tumor–immune cell interactions across disease sites^15–18^. These advances have come from focused characterization of the non-malignant components of the TME; however, the malignant epithelial subsets have been defined and annotated inconsistently, with no shared classification across cohorts^19,20^. Given the extensive chromosomal rearrangements and paucity of recurrent somatic mutations in HGSC^3,4^, HRD status remains the only molecular characteristic used to stratify the disease^3,4^. No unified framework describes epithelial diversity in HGSC, explains how it arises within and across tumors, or reconciles the states defined in individual studies.

In HGSC, epithelial state has been linked to the surrounding tissue, but which environmental features pattern that state, and how consistently they act across tumors, remain unresolved. Vázquez-García et al. demonstrated that epithelial signaling programs vary independently of clonal identity within individual patients and recur across patients with different genotypes^11^, suggesting that recurrent epithelial heterogeneity reflects environmentally programmed plasticity. Similar findings exist in other cancers: pancreatic cancer organoids collapse to a common state in culture irrespective of genotype and adopt distinct states under defined microenvironmental conditions^21^; in glioblastoma, malignant states are spatially organized around microenvironmental features such as hypoxic gradients^22^. HGSC tumors are structurally heterogeneous, exhibiting both solid and papillary growth patterns, as well as detached spheroids within peritoneal ascites, each defining a distinct microenvironment capable of shaping epithelial states. Resolving these recurring malignant states alongside the environmental features that pattern them would establish a unified framework for HGSC heterogeneity.

Here we examine how the local environment shapes epithelial state across HGSC. We integrated scRNA-seq data from 13 published studies into an atlas of approximately 2 million cells from 362 samples and generated targeted spatial transcriptomics of 8 whole tissues and tissue microarrays representing 378 patients. We identify a dominant axis of secretory epithelial polarization, from progenitor-like SecA to injury-associated SecB, that recurs across tumors and varies within genetic clones. In tumors, SecB polarization tracks with distance from vasculature, in hypoxic regions where HIF/NF-κB-driven survival and glycolysis replace mitogenic signaling. SecB cells are enriched in ascites. Lymphocytes are excluded from these regions, whereas macrophages adopt a hypoxic, glycolytic program that mirrors epithelial adaptation. These macrophages co-organize with SecB cells into an immune-protected niche poised for dissemination. The SecB state is largely lost in patient-derived organoids and only partially recovered by environmental inputs such as IFNγ, and it is incompletely recapitulated with *in vivo* models. Clinically, SecB polarization predicts worse survival after standard chemotherapy across independent cohorts, yet it identifies the patients who benefitted from bevacizumab in the ICON7 trial^23^. Together, these findings link HGSC tumor architecture to epithelial state, the immune niche, and treatment response.

## RESULTS

### Secretory epithelial cells adopt conserved polarized states in HGSC

To resolve the cellular landscape and transcriptional diversity of HGSC, we integrated scRNA-seq datasets from 13 independent studies (**Figure 1a,b**, **Figure S1**, **S2**, **Table S1**). This atlas comprises 1,980,703 cells from 158 patients across 362 tissue samples spanning 12 anatomic sites: primary tumors (adnexa), common metastatic sites (omentum, peritoneum, bowel), ascites, and rarer locations (lymph node, diaphragm, liver, upper quadrant, pelvic cavity, bladder, colon; **Figure 1c**, **Figure S3**). The atlas includes both treatment-naïve and post-treatment samples (**Figure 1d**, **Figure S3**). We adopted a hierarchical annotation scheme, identifying 13 broad cell type lineages that we further resolved into 51 discrete cell types and states (**Figure 1e, Figure S4, Table S2, S3**).

**Figure 1.**
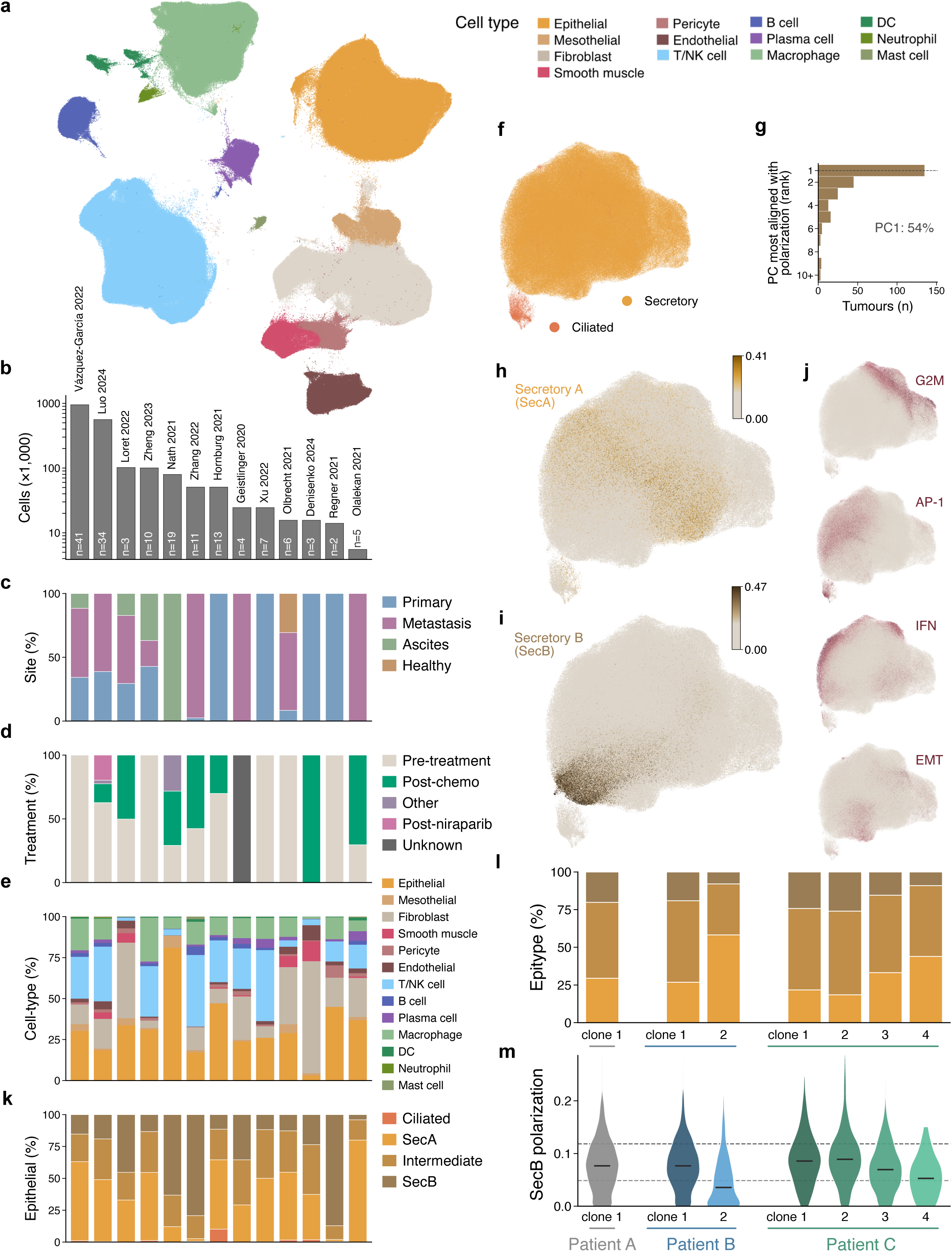
An integrated single-cell atlas identifies a conserved axis of secretory epithelial cell polarization in high-grade serous ovarian carcinoma. (**a**) UMAP of 1,980,703 cells from 13 integrated scRNA-seq studies colored by cell type annotation (13 cell types). (**b**) Bar graph representing cell counts per dataset with bars annotated with patient counts (n) per study (log scale). (**c–e**) Proportional stacked bar graphs demonstrating the per-study (**c**) anatomic site composition grouped by primary (adnexa), metastatic (non-primary site), ascites or healthy samples; (**d**) treatment status composition; and (**e**) cell type composition. (**f**) UMAP of the integrated epithelial compartment (575,366 cells) with a large dominant secretory population (golden) and small ciliated population (orange). (**g**) Bar graph showing the number of tumors within each principal component rank, most correlated with polarization score. (**h,i**) Epithelial compartment UMAP with NMF factor usage visualized by color gradient for (**h**) SecA or (**i**) SecB. (**j**) Epithelial compartment UMAP visualizing additional NMF factors, including G2M / cell cycle, early-stress/ AP-1, IFN signaling, and EMT. (**k**) Proportion of SecA, Intermediate, SecB or ciliated epithelial cells per study. (**l**) Stacked bar graph showing the secretory epithelial state composition of three representative tumors (one monoclonal, two polyclonal) inferred using CopyKAT across HGSC atlas samples. (**m**) Violin plot visualizing the distribution of SecB polarization scores across the CopyKAT clones represented in (**l**), with dashed lines representing the atlas thresholds for Intermediate (lower dashed lined) or SecB (upper dashed line) polarization score.

The integrated epithelial compartment (575,366 cells) included a small ciliated subset (6,678 cells, 1.2%) defined by canonical fallopian tube ciliated cell markers (*FOXJ1, CAPS, RSPH1, TPPP3*), with the remainder representing a dominant secretory population (*PAX8, WFDC2*; **Figure 1f**, **Table S4**). Within the secretory cell subset, markers associated with lineage-related differentiation were not uniformly expressed. To resolve coordinated transcriptional programs within the secretory compartment, we applied non-negative matrix factorization, which identified two programs that represent an axis of variation among malignant secretory cells (**Figure S5, Table S5**). Importantly, this axis represents the dominant coordinated transcriptional program across tumors (principal component (PC)1 in 54% of samples, PC1-3 in 82%; **Figure 1g**). One pole, we termed Secretory A (SecA), was enriched for developmental transcription factors and progenitor markers: *MECOM, SOX17, FBXO21, PBX1, LGR5, LPAR3,* and *RCN2* (**Figure 1h**, **Figure S6**). The other pole, which we termed Secretory B (SecB) was enriched for keratins and mucosal differentiation genes: *KRT17, LCN2, TACSTD2, KRT7, KRT19, SLPI,* and *PRSS22* (**Figure 1i**, **Figure S6**). This axis was independent of programs defining cell cycle, stress response, interferon, EMT, or metallothionein signatures (**Figure 1j**, **Figure S5**). To facilitate downstream comparisons, we partitioned cells along this axis into three groups: SecA, Intermediate, and SecB, based on their factor usage. These boundaries reflect divisions along a continuum rather than distinct cell identities (**Figure 1k**).

Having established the axis is independent of cell cycle and stress programs, we next asked whether it arises from genetic heterogeneity, leveraging inferred copy number variation (CNV) profiles. Within individual clones, we observed cells at both the SecA and SecB poles of the secretory axis (**Figure 1l,m, Figure S7**). This suggests that these states are not strictly driven by clonal divergence, though individual clones may differ in their propensity to activate specific programs.

### SecB polarized cells represent a quiescent state, enriched in ascites

SecA and SecB cells exhibit distinct regulatory and metabolic features (**Figure 2a**). We characterized the transcriptional programs of SecA and SecB cells through gene set enrichment (Hallmark), and inferred pathway (PROGENy), transcription factor (DoRothEA) and metabolic activity (scFEA). SecA cells were enriched for proliferation and growth programs, including MAPK signaling (PROGENy); MYC and E2F targets (Hallmark, DoRothEA); and metabolic flux estimation consistent with biosynthetic growth (**Figure 2a**, **Figure S8**). In contrast, SecB cells were enriched for signals related to environmental adaptation, including hypoxia, p53, VEGF, and TGFβ, and were metabolically more glycolytic (**Figure 2a**). Cell cycle analysis confirmed this distinction: SecA cells had the highest proliferating fraction (37.8% S/G2M), whereas SecB cells were predominantly in G1 (80.6% G1, **Figure 2b**).

**Figure 2.**
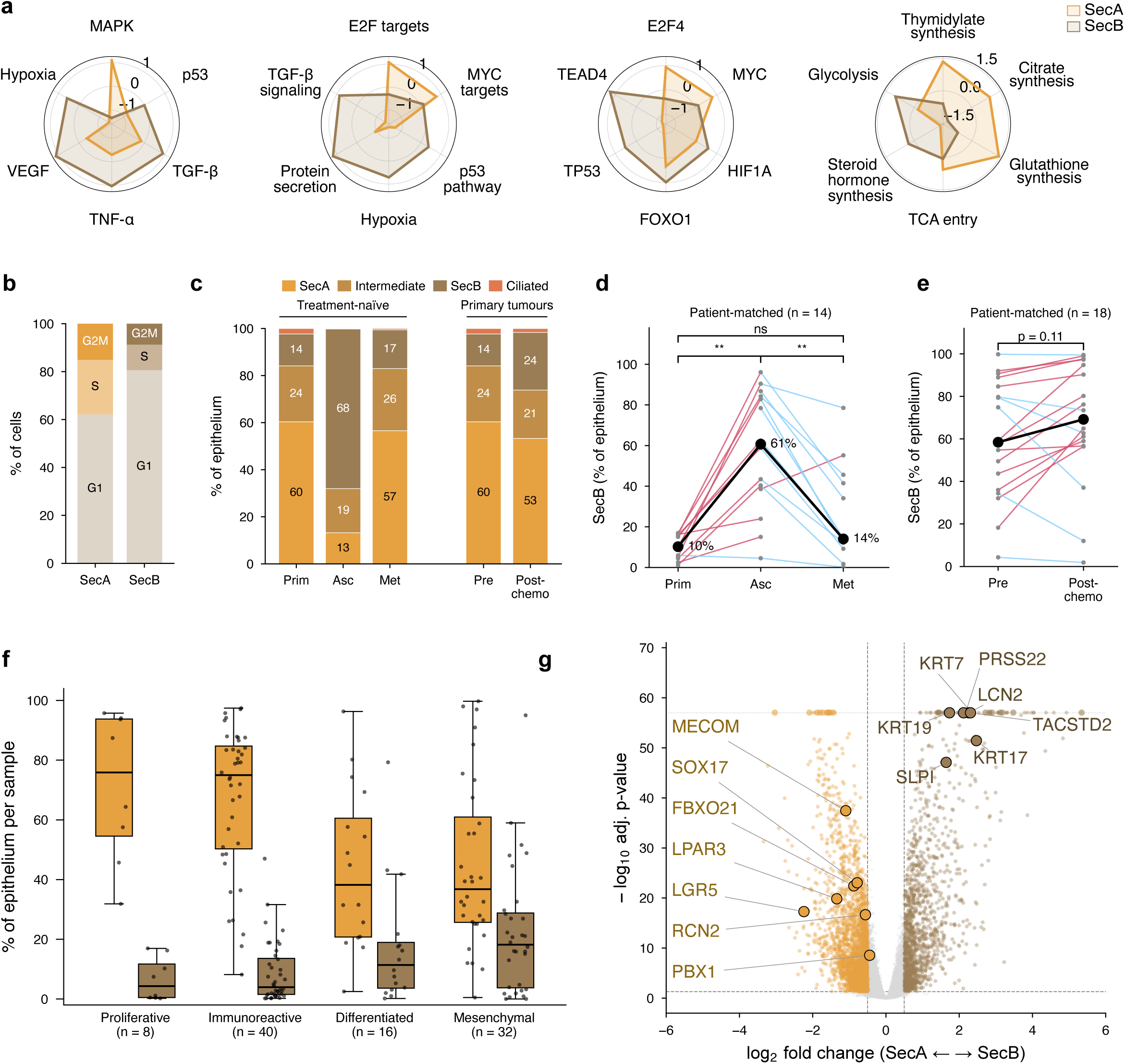
Transcriptional programs, cell cycle states, and distribution of secretory epithelium across anatomic sites and treatment conditions. (**a**) Radar plots of inferred pathway activity (PROGENy), gene set enrichment (Hallmark), transcription factor activity (DoRothEA) and metabolic flux estimation (scFEA) for SecA and SecB epithelium. (**b**) Cell cycle phase distribution for SecA and SecB. (**c**) Composition of epithelial compartment by anatomic site (treatment naïve, primary n = 99 samples from 81 patients; ascites n = 40 samples from 40 patients; metastases n = 131 samples from 89 patients) and composition of primary tumors by treatment status (pre-treatment n = 99 samples from 81 patients; post-chemotherapy n = 12 samples from 12 patients). (**d**) Patient-matched SecB proportion across anatomic sites (n = 14) and (**e**) across treatment conditions (n = 18, not site-matched). Matched patient samples were statistically compared by two-sided Wilcoxon signed-rank test (**p <0.01). (**f**) Box plot representing the proportion of SecA or SecB epithelium within pre-treatment primary tumors that were scored by consensusOV molecular subtype (n = 96 samples from 77 patients). (**g**) Volcano plot of genes differentially expressed between SecA and SecB epithelium, computed on pseudobulk profiles paired within tumors (n = 522 profiles from 261 tumors; DESeq2, ∼tumor + state; Benjamini–Hochberg FDR). Markers for SecA (*SOX17, LPAR3, LGR5, FBXO21, MECOM, RCN2, PBX1*) and SecB (*KRT7, KRT17, KRT19, SLPI, LCN2, TACSTD2, PRSS22*) used to score polarization of epithelium downstream are highlighted.

We quantified the proportion of each epithelial state across anatomic sites and after chemotherapy (**Figure S9**). Among pre-treatment samples, ascites was strongly enriched for SecB cells (68%), whereas primary and metastatic tumors showed similar SecB proportions (14% and 17%, respectively); this held in a subset of patient-matched multi-site samples (n = 14, median 60.7% in ascites vs 10.2% in primary; **Figure 2c**,**d**). In post-chemotherapy primary tumors, SecA proportions declined (from 60% to 53%) and SecB proportions increased (from 14% to 24%, **Figure 2c**). In a subset of patient-matched pre- and post-chemotherapy samples (n = 18), SecB increased after chemotherapy in 12 of the 18 patients (58% to 69%, p = 0.11, **Figure 2e**).

These findings suggest SecB cells may be dependent on the tumor microenvironment they exist within. We therefore asked if SecB cells are differentially enriched within the previously established molecular subtypes of HGSC tumors^3,4^. We categorized primary samples from the atlas into their TCGA-defined subtype using consensusOV, then explored the proportion of epithelial states within each (**Figure 2f**). We observed that SecA cells predominate in proliferative and immunoreactive tumors (median 76% and 75%) but were lower in differentiated and mesenchymal tumors (median 38% and 37%), while SecB was most abundant in mesenchymal tumors (**Figure 2f**). Together, these data suggest SecB polarization reflects a quiescent, metabolically adapted state that may confer an advantage under environmental stress, such as hypoxia and after exposure to chemotherapy. To explore secretory polarization across independent models and platforms, we derived a SecA/SecB gene signature from each NMF factor loading and used it to score polarization in downstream analyses, providing consistent, atlas-aligned thresholds (**Figure 2g**, **Table S6**).

### In vitro and in vivo models of HGSC incompletely recapitulate secretory polarization

To assess whether we could model and perturb this secretory polarization *in vitro*, we performed snRNA-seq on eight patient-derived organoid (PDO) models derived from ascites (n = 4) or primary tumors (n = 4) (**Figure 3a**). Compared to the atlas, PDO models were strongly skewed toward SecA, with fewer than 1% of cells reaching the SecB signature threshold (**Figure 3b**). PDOs established from ascites samples, however, consistently exhibited higher SecB scores than those from primary tumors (p = 0.029), suggesting partial retention of this state from the ascites (**Figure 3b**).

**Figure 3.**
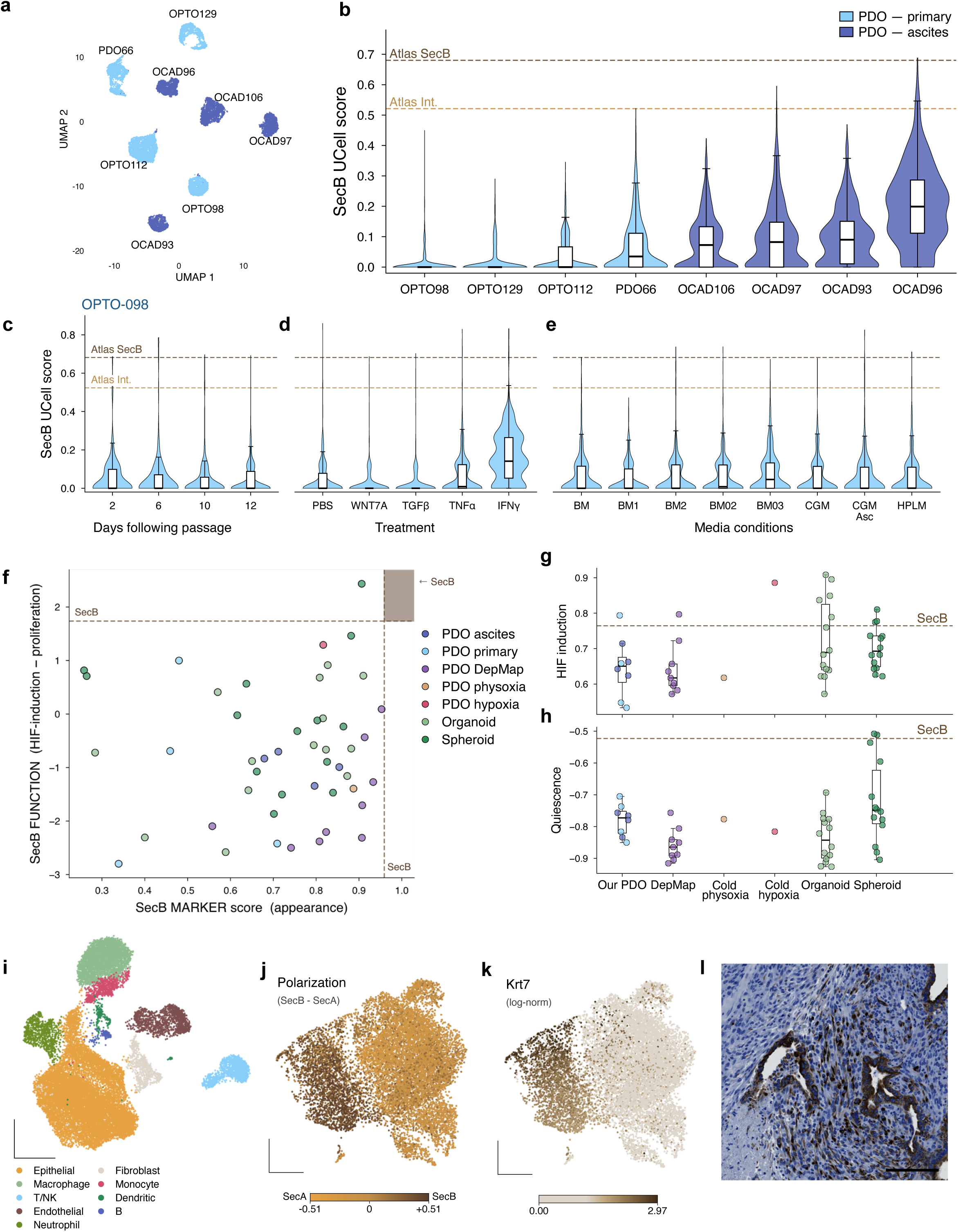
Secretory polarization in patient-derived organoids (PDOs) and a syngeneic fallopian-tube-derived model of HGSC. (**a**) UMAP of snRNA-seq from eight PDO models of HGSC derived from primary tumor (light blue) or ascites (dark blue). (**b**) SecB UCell score distributions per PDO model with dashed lines indicating atlas median SecB UCell scores for Intermediate and SecB epithelium. Ascites and primary-derived models were compared by two-sided Mann–Whitney test, n = 4 vs 4, p = 0.029. (**c–e**) SecB UCell score distributions of OPTO98 (**c**) over time in standard Matrigel culture conditions (day 2**–**12), (**d**) following treatment with microenvironmental factors including PBS control, WNT7A, TGFβ, TNFα, and IFNγ for 6 days, or (**e**) in a variety of media conditions including basal media (BM), complete growth media (CGM), and human plasma-like medium (HPLM) for 6 days. (**f**) Scatter plot quantifying the expression of SecB markers (*KRT17*, *LCN2*, *TACSTD2*, *KRT7*, *KRT19*, *SLPI*, *PRSS22*) against functional score, which was defined as a z-score of HIF induction minus proliferation in various HGSC *in vitro* model systems, including our PDOs, DepMap PDOs, cold-co-culture physioxia PDOs, and cold-co-culture hypoxia PDOs, in addition to matched organoid versus spheroid models. (**g, h**) The two functional scores to model the SecB state, which quantify (**g**) HIF induction and (**h**) quiescence (defined as negative proliferation) for each model, grouped by dataset. (**i**) UMAP of scRNA-seq from the KPCA.B orthotopic syngeneic mouse model of HGSC, colored by cell type annotation. (**j,k**) UMAP of the epithelial compartment (n = 12,429 cells), colored by (**j**) secretory polarization score and (**k**) expression of *Krt7*, a SecB marker well conserved from human to mouse. (**l**) Representative KRT7 immunohistochemistry, showing KRT7 secretory cells lining luminal regions away from the stroma; day-26 tumor, scale bar represents 100 µm.

Consistent with environmental maintenance of SecB, polarization scores in the OPTO98 PDO declined progressively over time in standard Matrigel-embedded, glucose-rich culture (**Figure 3c**). To attempt to identify the signals responsible, we treated organoids with candidate microenvironmental factors (IFNγ, TNFα, TGFβ, and WNT7A). Of all these perturbations, only IFNγ induced SecB gene expression across the population (**Figure 3d**), suggesting a role for the immune microenvironment in initiating or maintaining secretory polarization. We additionally tested if we could induce SecB polarization by differing culturing medias, but unfortunately no medium was able to recapitulate the SecB polarization observed in the atlas (**Figure 3e**).

We next asked whether the SecB state was represented across publicly available RNA-seq datasets of organoids and spheroids derived from HGSC (**Figure 3f**). We scored these models on their ability to recapitulate the SecB markers and function (HIF induction and quiescence) against the atlas reference (**Figure 3f–h**). While models cultured in hypoxic conditions recapitulate HIF induction (**Figure 3g**), proliferation rates remain elevated across models (**Figure 3h**). These data suggest that our current organoid and spheroid models lack the diverse cellular interactions and tissue-level features required to recapitulate the SecB state.

To explore the possibility of modeling SecB *in vivo*, we profiled the cellular composition of one primary endpoint tumor from the KPCA.B orthotopic syngeneic model of HGSC by scRNA-seq (**Figure 3i**). We identified several programs driving secretory cell heterogeneity within the KPCA.B model (**Figure 3j, Figure S10**), with expression of the top conserved SecB marker, *KRT7*, establishing a clear axis (**Figure 3k**). Immunohistochemistry on KPCA.B tumors, localized KRT7+ cells to luminal regions, where they appeared distant from the stroma and were frequently adjacent to immune cells (**Figure 3l**). Although the syngeneic model may be able to partially model a SecB-like state, no resulting NMF program reflected the human SecB polarization (**Figure S10**). The species-driven differences in epithelial and immune cell regulation and divergence from the disease’s natural progression by orthotopic cell injection may limit its ability to reflect the human SecB state.

### Tumor architecture establishes a metabolic gradient aligned with secretory polarization

To identify the environmental features that drive the SecB state, we designed a custom targeted spatial transcriptomics panel (10x Genomics Xenium) to resolve cell types, epithelial programs, signaling pathways, immune function, therapeutic targets, and clinical classifiers relevant to HGSC (**Table S7**). We applied this panel to 8 whole HGSC tumors (n = 4 treatment-naïve, n = 4 post-chemotherapy, approximately 1.9 million cells, **Table S8**), to two tissue microarrays (TMAs) of primary HGSC representing a combined 378 patients (503 cores, 1.5 million cells: TMA A, 95 patients, 82 profiled in duplicate cores; TMA B, 283 patients), and to 15 healthy fallopian tube epithelial (FTE) tissue cores (34,281 cells, **Figure 4a**, **Figure S11–13**). Eighteen cell types, including ciliated and secretory epithelial cell states, were annotated by SingleR using the scRNA-seq atlas as a qualitative reference (**Figure S14**). Hierarchical clustering revealed broad diversity in HGSC tissue composition, with whole tissues spanning the range observed across the TMA patients (**Figure 4a**). Secretory cells were scored for polarization (SecB – SecA) and annotated according to atlas-aligned thresholds, with polarization scoring significantly higher in the HGSC secretory cells compared to those within the healthy FTE tissue (per-unit medians, p = 0.003, **Figure S15**).

**Figure 4.**
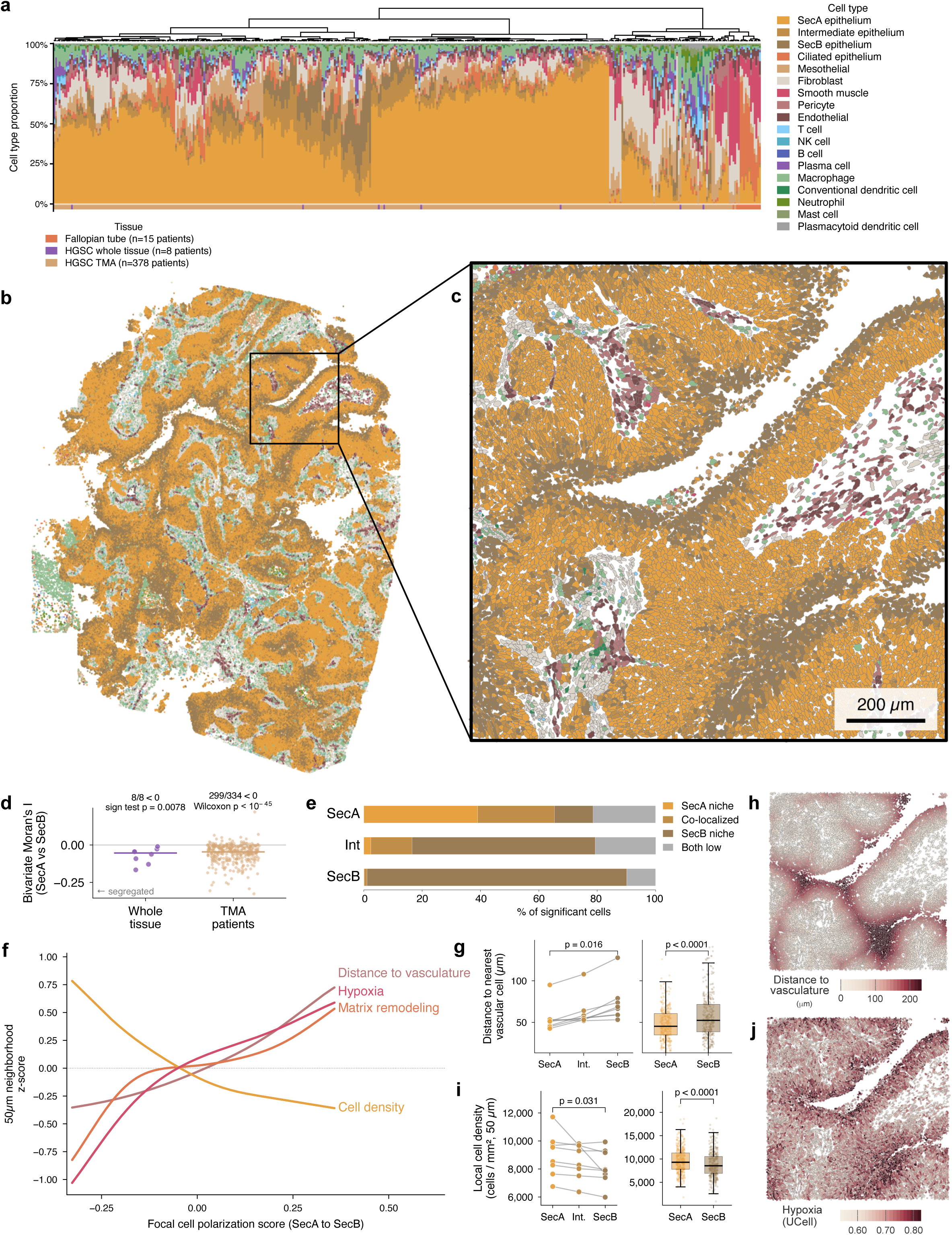
SecB-polarized secretory cells co-localize within avascular and hypoxic niches. (**a**) Per-sample cell-type composition across the Xenium cohorts: fallopian tube epithelium (15 cores), whole HGSC tissues (8 sections), and HGSC TMAs (378 patients with ≥500 cells; TMA A and TMA B). Samples are ordered by hierarchical clustering, and the bottom row indicates tissue type. (**b**) Whole-tissue cell-type map of sample WT01. (**c**) ROI from sample WT01 indicated by the ROI box on the whole-tissue map in (**b**), colored by cell type. (**d**) Global bivariate Moran’s I between each secretory cell’s SecA UCell score and its neighbors’ SecB UCell scores (50 µm neighborhoods) per whole tissue (n = 8) and per TMA patient (mean over cores with ≥500 secretory cells; n = 334). Negative values indicate segregation; lines show medians. Whole tissue: exact sign test; TMA: one-sample Wilcoxon signed-rank test. (**e**) Stacked bar graph showing the percentage of each secretory cell when assigned to locally defined (50 µm) spatial neighborhoods (SecA neighborhoods, SecB neighborhoods, both or neither) by bivariate local indicators of spatial association (BiLISA). (**f**) Generalized additive models of distance to vasculature, cell density (number of neighbors per neighborhood, cells/mm^2^), and UCell pathway scores (z-scored) for matrix remodeling (*ADAM10, ADAM17, MMP7, F3, SERPINE1, MMP11*) and hypoxia (*CAT, ENO1, EPAS1, HIF1A, LDHA, NFE2L2, PDK1, PGK1, PRDX1, SLC2A1, TXN, VEGFA)* as a function of focal cell polarization score across all secretory cells in the whole tissue samples. (**g**) Distance to nearest vascular cell (pericyte or endothelial cell) measured by Euclidean distance for SecA, Intermediate or SecB cells within each whole-tissue (left; paired Wilcoxon, p = 0.016) and SecA and SecB cells per TMA sample (right; n = 361, p < 0.0001). (**h**) Spatial plot of ROI from sample WT01 in (**b**) visualizing the distance-to-vasculature gradient. (**i**) Local cell density (all cells within 50 µm, cells/mm^2^; median per sample) around SecA, Intermediate and SecB cells in each whole tissue (left; n = 8; paired Wilcoxon, p = 0.031) and SecA and SecB cells per TMA patient (right; n = 361; p < 0.0001). (**j**) Spatial plot of ROI from sample WT01 visualizing the UCell-scored hypoxic gradient.

Across all eight whole tissues, SecA and SecB scores were spatially segregated (**Figure 4b,c)**: a cell’s SecA score was negatively associated with its neighbors’ SecB scores within each whole tissue (bivariate Moran’s I −0.01 to −0.17; sign test p = 0.008) and in 299 of 334 TMA patients (Wilcoxon p < 0.001), forming continuous spatial polarization gradient toward the lumen of papillary growths (**Figure 4d**). Whereas SecA cells were broadly distributed throughout HGSC tissue, SecB cells were spatially concentrated: 89% of SecB cells assigned to a neighborhood resided in neighborhoods enriched for other SecB cells, suggesting that this adaptation occurs in discrete, spatially coherent niches (**Figure 4e**). To understand what shapes these niches, we characterized the local microenvironment (50 µm radius) of each secretory cell (1,235,350 cells) and quantified regional cell type composition, bulk environmental signaling, and cell-specific gene expression. We used generalized additive modeling to map local environmental changes along the polarization axis (pathway UCell scores using custom gene sets, **Table S9**). In the tissue maps, SecB cells lay toward the lumen of papillary structures (**Figure 4b,c**), and they were enriched with increasing distance from the vasculature (**Figure 4f– h, Figure S16**). These shifts coincide with reduced local cell density (**Figure 4f,i**) and increased expression of matrix remodeling genes (e.g. *ADAM17, MMP7, SERPINE1,* **Figure 4f**). These architectural and compositional changes are associated with shaping the local metabolic environment; as secretory cells move away from blood vessels, expression of hypoxia-associated genes (e.g. *ENO1*, *HIF1A*, *VEGFA*) increased (**Figure 4f,j**).

### SecB cells and macrophages adapt to hypoxic niches through a shared glycolytic survival program

We next examined how transcriptional programs within the secretory cells adapt along the secretory polarization axis (**Figure 5a–d**). Within secretory epithelial cells, we observed a consistent decline across markers of proliferation and growth as cells adapt toward SecB, including: proliferation signals (*MKI67*, *PCNA*, *TOP2A*), oncogenic growth signaling pathways (Myc [*MYC, MAX, MYCL*], PI3K [*PIK3CA, AKT1*, *MTOR*], Wnt [*FZD3*, *CTNNB1*, *TCF7L2*], and Notch [*JAG1*, *NOTCH1*, *NOTCH2*]), DNA damage repair genes (*BRCA1/2*, *ATM*, *CHEK2*), and epigenetic remodeling signals (*DNMT3A, HDAC1, KDM5A*) (**Figure 5a,c,d; Figure S17**). Instead, SecB cells maintain survival through increased *TNFA* expression, NF-κB signaling (*BIRC3*, *ICAM1*, *NFKBIA*), and HIF-driven glycolysis (*SLC2A1*, *LDHA*, *SLC16A3*; **Figure 5a,c,d**). Consistent with reduced proliferation and growth signaling, SecB cells had significantly smaller nuclei than SecA cells across our Xenium tissues (median 39.5 vs 47.7 µm², p = 0.008) and across the TMA cohort (n = 361 patients, p < 0.001; **Figure 5b**). These signals suggest that SecB cells adapt to their hypoxic environment by trading proliferation for stress-tolerant survival.

**Figure 5.**
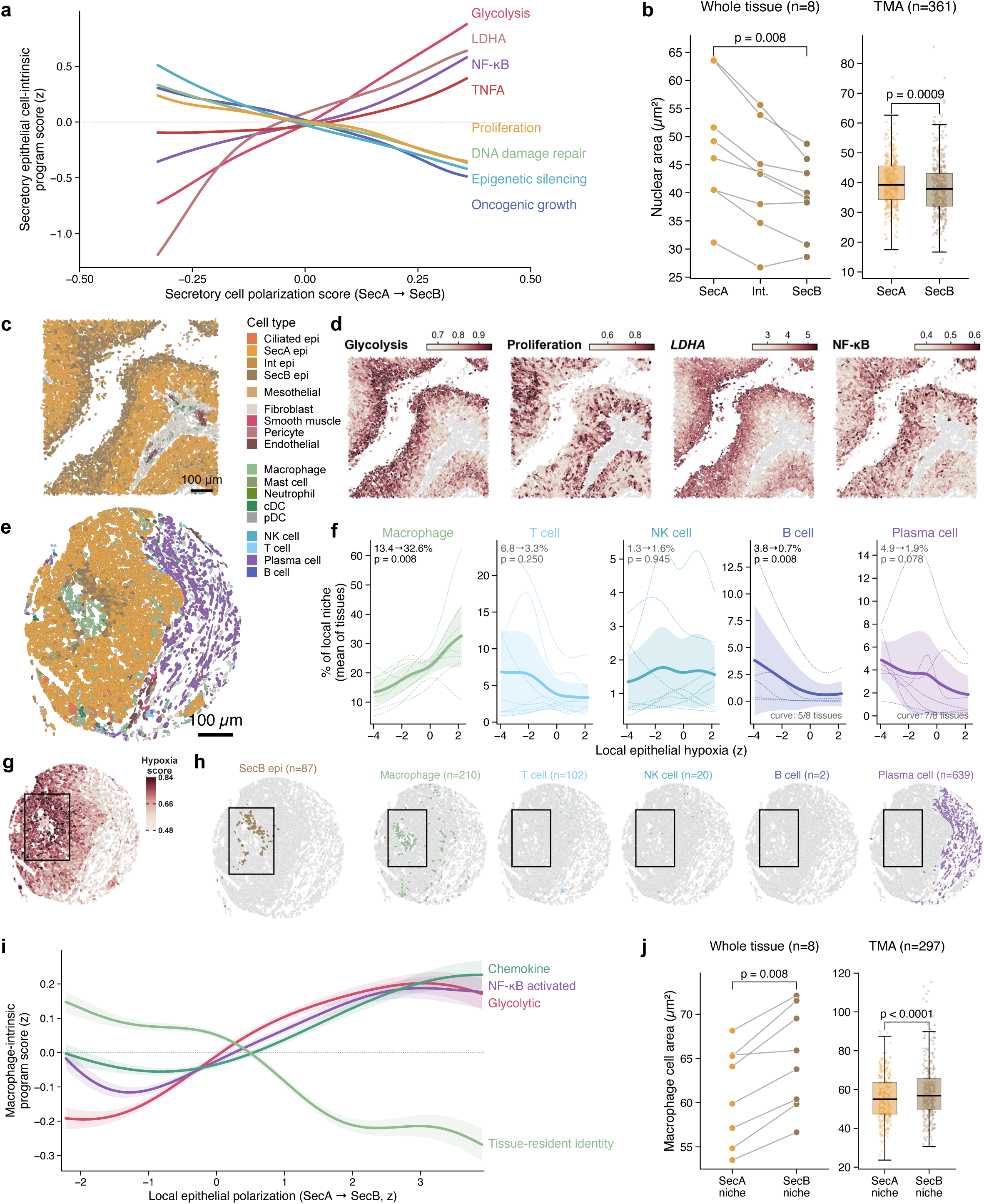
SecB cells and macrophages adapt to hypoxic niches through a shared glycolytic survival program. (**a**) Generalized additive models of epithelial-specific UCell pathway scores (z-scored) plotted as a function of focal cell polarization score across eight whole-tissue samples. (**b**) Nuclear area (µm²) of SecA, Intermediate and SecB cells across whole tissue (left, n = 8) and SecA and SecB cells per TMA patients (right, n = 361). (**c**) Spatial map of a SecB-enriched ROI from whole-tissue sample WT01, colored by cell type annotation, and (**d**) the same ROI visualizing (left to right) UCell pathway scores for glycolysis, proliferation, LDHA expression, and NF-κB pathway expression in secretory epithelial cells (other cells grey). (**e**) Representative TMA core (TMA A, TMA-P003) colored by cell type annotation. (**f**) Immune cell proportion of the local niche (percentage of non-epithelial cells within 50 µm) as a function of local epithelial hypoxia (z-scored) for (left to right) macrophages, T cells, NK cells, B cells, and plasma cells across whole-tissue samples (n = 8). Dotted lines represent the individual tissue fits behind the pooled curve (B cell and plasma cell curves average 5 and 7 tissues, respectively). (**g**) Hypoxia UCell score across the representative TMA core; the box highlights the SecB-enriched hypoxic region. (**h**) The representative TMA core, highlighting (left to right) SecB cells, macrophages, T cells, NK cells, B cells, and plasma cells. (**i**) Generalized additive models of macrophage-intrinsic UCell program scores (z-scored) as a function of local epithelial polarization, including for glycolysis (*LDHA*, *ENO1*, *PGK1*, *ALDOA*), chemokine production (*CXCL10*, *CXCL11*), NF-κB activation (*ICAM1*, *NFKBIA*, *BIRC3*), and tissue-resident identity (*A2M, C1QC*). (**j**) Median macrophage cell area in SecA compared to SecB niches (lowest versus highest tertile of local epithelial polarization within each unit) across whole tissue (left, n = 8) and TMA patients (right, n = 297). Statistical comparisons between samples were performed by two-sided paired Wilcoxon signed-rank test.

The hypoxic niches that enriched for SecB cells also showed coordinated shifts in immune composition. By modeling the composition of each cell’s neighborhood as a function of local epithelial hypoxia across the eight whole tissues, we found that macrophages and B cells followed opposite trajectories: macrophages rose from 13.4% to 32.6% of non-epithelial cells in the most hypoxic niches, whereas B cells fell from 3.8% to 0.7% (both p = 0.008, 8 of 8 tissues; **Figure 5e–h**). Within hypoxic regions, T cells and plasma cells trended lower, and NK cells were unchanged (**Figure 5e-h**). Macrophages not only persisted in the most metabolically stressed niches but activated a transcriptional program mirroring the SecB survival program (**Figure 5i,j; Figure S18**). Macrophages in SecB regions adopted glycolytic (*LDHA*, *ENO1*, *PGK1*, *ALDOA*) and NF-κB (*ICAM1, NFKBIA, BIRC3*) programs with reduced tissue-resident identity (*A2M*, *C1QC*, **Figure 5i**). They concurrently maintained chemokine production (*CXCL10*, *CXCL11*), signaling for lymphocyte recruitment into the metabolically hostile niche (**Figure 5i**). This transcriptional program also aligned with a coordinated morphological shift, with SecB-adjacent macrophages being larger than SecA-adjacent macrophages (**Figure 5j**).

### SecB epithelium co-opts a hypoxic macrophage state into a self-sustaining pro-dissemination niche

To better describe the relationship between macrophages and SecB cells sequestered within the niche, we performed Visium HD on one of our Xenium tissues (WT01, **Figure 4**). This sample was selected as it held the highest proportion of SecB (**Figure 4a**, **Figure 6a Table S8**). By Visium HD, we were able to further confirm that the spatially driven SecB polarization aligns with transcriptional expression of the hypoxia-associated *CA9* (**Figure 6b–d**). We compared the macrophage bins adjacent to SecB-high epithelium to distal macrophage bins and found that SecB macrophages upregulate expression of osteopontin (*SPP1*) (**Figure 6e**) and *MMP9*, among other known hypoxia response genes (**Table S10**). This transcriptional program aligned to the hypoxic macrophages within the atlas (**Figure 6f**) and reflected the glycolytic program we described (**Figure 5i**). Across our Xenium tissues, macrophages defined by this hypoxic program had twice the proportion of SecB neighbors among their epithelial neighbors compared to resident-like macrophages (**Figure 6g**).

**Figure 6.**
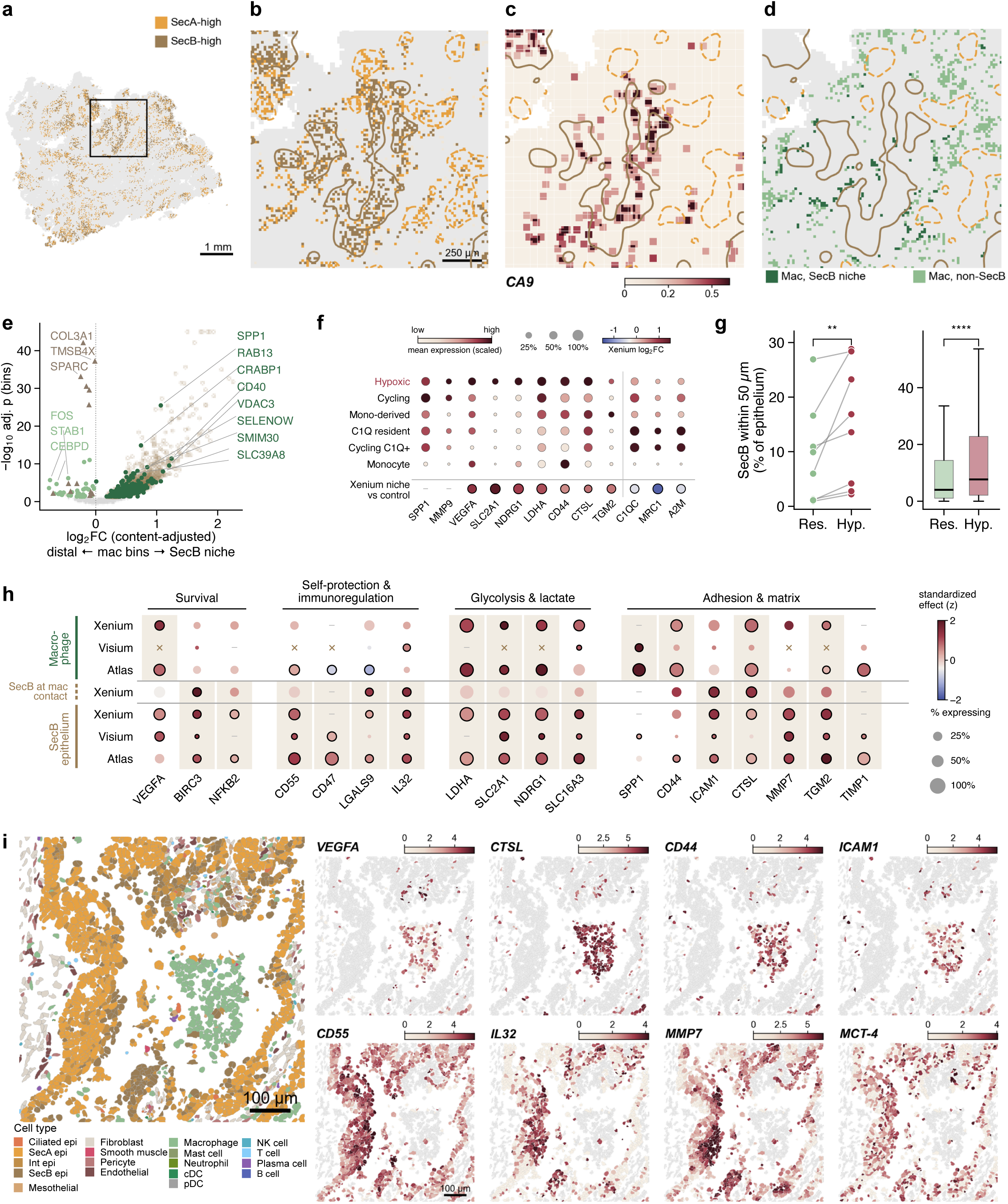
SecB epithelium co-opts a hypoxic macrophage state into a self-sustaining pro-dissemination niche. (**a**) Visium HD whole-transcriptome spatial map (16 µm bins) of whole-tissue sample WT01, colored by SecA-high and SecB-high epithelial bins (bottom and top tertiles of the SecB–SecA polarization score among epithelial bins); boxed region is magnified in (**b–d**). (**b**) SecA-high and SecB-high epithelial bins within the region of interest, with SecB and SecA territories outlined. (**c**) CA9 expression (log1p CP10k, 3 × 3-bin mean) within the region of interest, overlaid with SecB and SecA territories. (**d**) Macrophage bins within the region of interest classified as SecB-niche (≥2 SecB-high bins within 48 µm, dark green) or non-SecB (light green), overlaid with SecB and SecA territories. (**e**) Volcano plot of genes differentially expressed between SecB-niche and distal macrophage bins in sample WT01 (log2 fold change adjusted for macrophage content; bin-level Mann–Whitney test, Benjamini–Hochberg FDR). Macrophage-derived genes (dark green, up; light green, down) are highlighted; transcripts attributable to adjacent SecB epithelium (open circles) or stroma (triangles) are shown but excluded. (**f**) Dot plot of macrophage subtypes in the HGSC atlas (n = 295,296 cells) showing scaled mean expression (color) and percentage of expressing cells (size) of hypoxic-macrophage and tissue-resident genes, with the corresponding log2 fold change in Xenium SecB-niche versus control macrophages (bottom row, n = 8 whole tissues; outlined dots p < 0.05). (**g**) Percentage of SecB cells among epithelium within 50 µm of resident-high compared to hypoxic-high macrophages (glycolytic-minus-resident tertiles) across whole tissue (left, n = 8) or per patient in the TMA (right, n = 311). (**h**) Cross-platform concordance of genes grouped by function (survival, self-protection and immunoregulation, glycolysis and lactate export, adhesion and matrix remodeling) in macrophages (top: Xenium SecB-niche versus control macrophages, n = 8; Visium HD SecB-niche versus distal macrophage bins, WT01; atlas hypoxic versus other macrophages, n = 181 paired samples) and SecB epithelium (bottom: Xenium SecB versus SecA cells, n = 8; Visium HD SecB-high versus SecA-high bins, WT01; atlas pseudobulk SecB versus SecA, n = 261 tumors). Color represents effect size standardized within each comparison, size represents percentage of expressing cells, and outline indicates statistical significance; × denotes transcripts attributable to adjacent SecB epithelium, and shading marks the cell type in which each gene is enriched. (**i**) Representative spatial plots of an ROI from sample WT02 (700 µm) visualizing (left) cell type annotation; (top row) macrophage expression of VEGFA, CTSL, *CD44* and *ICAM1;* and (bottom row) epithelial expression of *CD55*, *IL32*, *MMP7* and MCT-4 (*SLC16A3*). Non-focal cells are shown in grey, and color scales represent log1p CP10k (2nd–98th percentile). Statistical comparisons between samples were performed by two-sided Wilcoxon signed-rank test (**p < 0.01, ****p < 0.0001).

Within this niche, SecB cells and hypoxic macrophages exchange survival and immune-protective signals. We explored differentially expressed genes from SecB-niche versus control macrophages across our datasets, and within our Xenium we also looked at SecB cells directly in contact with (within 30µm of) adjacent a SecB-niche macrophages. Macrophages express *VEGFA*, while SecB cells in contact with them further upregulate their downstream *NF-κB* target *BIRC3* potentially amplifying their survival program (**Figure 6h,i**). SecB cells in turn protect themselves from macrophage phagocytosis and complement by upregulating *CD47* and *CD55*, respectively (**Figure 6h,i**). Additionally, we observed that SecB cells could be engaging in direct immunosuppression through the upregulation of *IL-32* and galectin-9 (*LGALS9*), with galectin-9 further increasing upon macrophage contact (**Figure 6h,i**). Notably, SecB cells upregulate *SLC16A3* (MCT-4), which likely contributes to the lymphocyte-hostile environment by acidifying the niche (**Figure 6h,i**).

This niche also contains a coordinated adhesion and matrix-remodeling program that could favor dissemination. *SPP1* receptor, *CD44,* was also found enriched on macrophages within the niche (**Figure 6h,i**). Additionally, SecB cells upregulated *ICAM1*, especially upon macrophage contact, which could provide SecB cells with a tether to immune or mesothelial cells (**Figure 6h,i**). These SecB cells and macrophages within this niche also likely collaborate to constantly remodel this environment. For example, SecB cells express *MMP7* and *TIMP1*, alongside macrophages that produce enzymes, including cathepsins such as *CTSL*, which can accelerate remodeling (**Figure 6h,i**). Both cells contribute to *TGM2* expression supporting local remodeling (**Figure 6h**). Together, these signals likely support homotypic and heterotypic adhesions within a niche marked by extracellular matrix remodeling (**Figure 6h,i**).

### Secretory polarization independently predicts survival and bevacizumab response

To assess the impact of secretory polarization on clinical outcomes, we performed survival analyses in patients represented by two primary, treatment-naive HGSC cores (OS n = 104, PFS n = 103). Patients in the highest polarization tertile (SecB-enriched) had worse 5-year OS than those in the lowest tertile (SecA-dominant; p = 0.011), and a trend towards worse 5-year PFS (p = 0.11; **Figure 7a,b**). In multivariable Cox regression adjusted for stage, residual disease and age, SecA proportion was protective for OS (HR = 0.68 per SD, 95% CI 0.51–0.90, q = 0.029) and PFS (HR = 0.79, 95% CI 0.63–0.98, q = 0.048), whereas SecB proportion was associated with worse OS (HR = 1.38, 95% CI 1.06–1.78, q = 0.030), with a similar trend for PFS (HR = 1.23, 95% CI 1.00–1.50, q = 0.051; **Figure 7c,d**). Consistent with this, the continuous per-patient polarization score (mean cell-level polarization across all secretory cells) was independently prognostic for OS (HR = 1.51, 95% CI 1.11–2.07, q = 0.029), with a similar trend for PFS (HR = 1.27, 95% CI 0.999–1.62, q = 0.051), indicating that the secretory polarization state, not just the abundance of either pole, carries clinical relevance (**Figure 7c,d; Table S11**).

**Figure 7.**
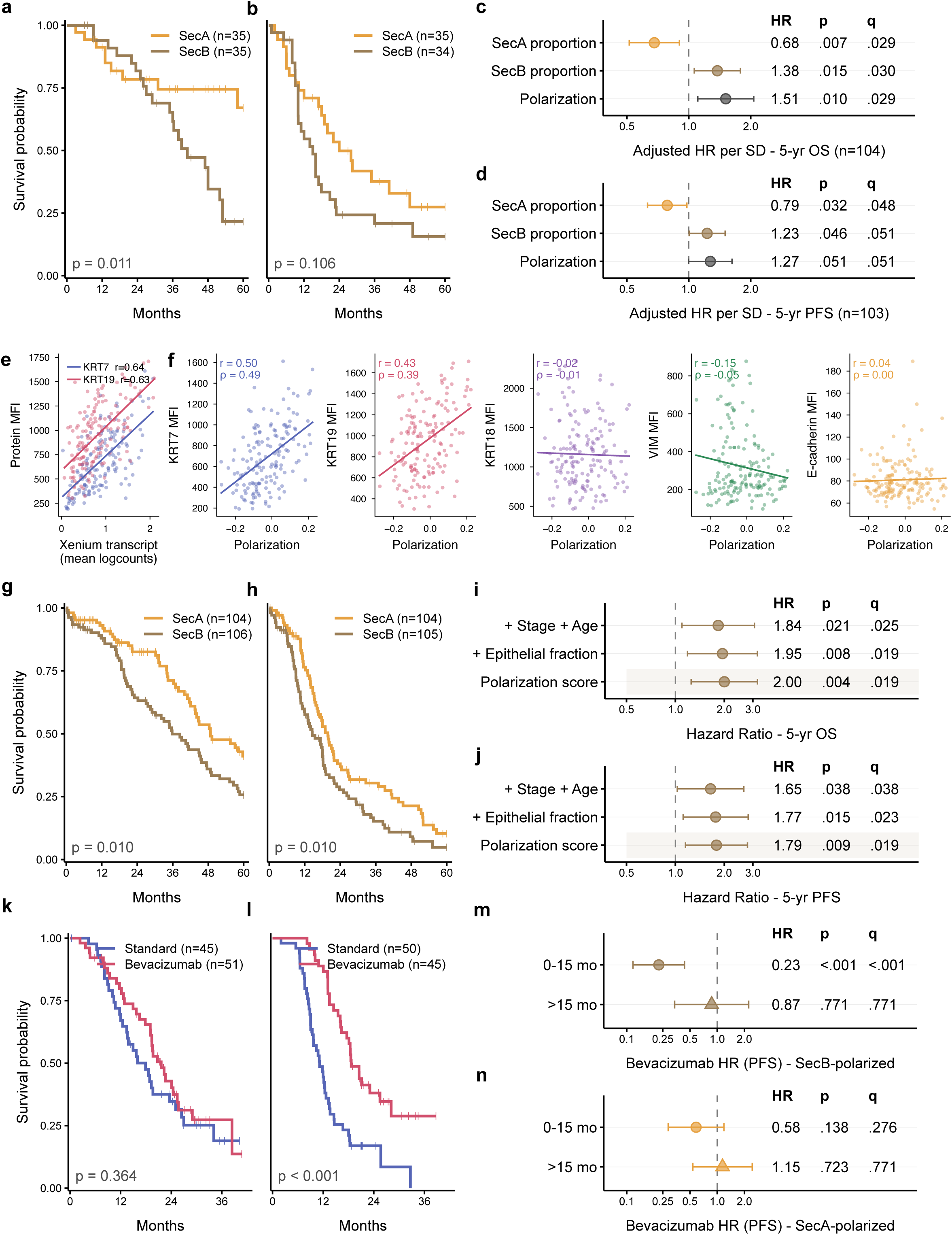
Secretory polarization independently predicts survival, is mirrored by KRT7 and KRT19 protein expression, and identifies bevacizumab-treatable tumors. (**a, b**) Kaplan–Meier curves for 5-year (**a**) overall survival (OS, n = 104; SecA-dominant = 35, SecB-enriched = 35) and (**b**) progression-free survival (PFS, n = 103; SecA-dominant = 35, SecB-enriched = 34) in patients with primary, treatment-naive HGSC represented by two tumor cores across our TMAs, comparing the lowest (SecA-dominant) and highest (SecB-enriched) tertiles of per-patient polarization score (middle tertile not shown); log-rank p shown. (**c**, **d**) Forest plots of multivariable Cox proportional hazards regression for 5-year (**c**) OS and (**d**) PFS of the SecA and SecB proportion of secretory cells and the mean cell-level polarization score (UCell), each model adjusted for stage, residual disease and age. Points show hazard ratios (HR) per 1 SD increase and whiskers 95% confidence intervals; p, Wald test; q, Benjamini–Hochberg FDR across the six tests shown. (**e**) Per-core correlation between Xenium mean transcript expression (mean log counts) and immunofluorescence (IF) protein mean fluorescence intensity (MFI) for KRT7 and KRT19 (n = 157 TMA cores). Pearson r shown. (**f**) Per-core correlation between mean polarization score (Xenium) and IF protein MFI for KRT7, KRT19, KRT18, VIM and E-cadherin. Pearson r and Spearman ρ shown. (**g, h**) Kaplan– Meier curves for 5-year (**g**) OS (SecA n = 104, SecB n = 106) and (**h**) PFS (SecA n = 104, SecB n = 105) in TCGA-OV, comparing the highest and lowest tertiles of polarization score. (**i, j**) Forest plots of Cox proportional hazards regression for the polarization score in the full TCGA-OV cohort for (**i**) OS and (**j**) PFS, shown unadjusted, adjusted for epithelial cell fraction, and additionally adjusted for stage and age ((OS n = 314, 311 with stage and age; PFS n = 312, 309). (**k, l**) Kaplan–Meier curves for PFS in FIGO stage III/IV patients from the ICON7 trial (GSE140082, n = 191), comparing carboplatin–paclitaxel alone (standard) with carboplatin–paclitaxel plus bevacizumab in (**k**) SecA-polarized tumors (n = 96; standard 45, bevacizumab 51) and (**l**) SecB-polarized tumors (n = 95; standard 50, bevacizumab 45), log-rank p shown; adjusted hazard ratios in **Table S12**. (**m, n**) Time-varying Cox proportional hazards regression of bevacizumab benefit (HR vs standard therapy for PFS) within (**m**) SecB-polarized and (**n**) SecA-polarized tumors, split at 15 months: 0–15 months (circles) or beyond 15 months (triangles), adjusted for FIGO stage, debulking status and age; q, Benjamini–Hochberg FDR across the four estimates shown.

We next correlated our Xenium transcript counts and polarization scores with the abundance of prognostic proteins previously measured on a subset of the same cores. We found that transcript counts and protein abundance were strongly correlated for SecB markers KRT7 and KRT19 (KRT7: Pearson r = 0.64; KRT19: Pearson r = 0.63; **Figure 7e**). Furthermore, mean polarization scores also correlated with protein abundance for these two SecB markers, while E-cadherin, VIM, and KRT18 showed no association with polarization score (**Figure 7f**). These results validate and contextualise earlier findings on this cohort demonstrating that KRT7 and KRT19 were independently associated with worse OS and PFS after correction for stage and residual disease^24^.

To validate these survival effects in an external cohort, we applied SecA-specific and SecB-specific gene signatures to bulk RNA-seq data from 314 TCGA-OV tumors, quantifying polarization as the difference between the z-scored SecB and SecA signatures^4^. Patients in the highest polarization tertile, SecB tumours, had significantly worse 5-year OS (p = 0.010) and PFS (p = 0.010) than those in the lowest tertile (**Figure 7g,h**). Cox proportional hazards regression confirmed that increased polarization was associated with worse OS (HR = 2.00, 95% CI 1.25–3.20, p = 0.004) and PFS (HR = 1.79, 95% CI 1.15–2.78, p = 0.009; **Figure 7i,j**). This association remained significant after adjustment for epithelial cell fraction, stage and age (OS HR = 1.84, 95% CI 1.10– 3.07, p = 0.021; PFS HR = 1.65, 95% CI 1.03–2.64, p = 0.038; **Figure 7i,j**).

Given that SecB niches are relatively avascular (**Figure 4g,h**), we next asked whether the degree of SecB polarization could influence response to the anti-VEGF antibody bevacizumab, which is routinely used to treat HGSC. To test this, we scored secretory polarization in pre-treatment samples from stage III/IV patients enrolled in the ICON7 trial (n = 191), and we classified them as SecA-polarized (n = 96) or SecB-polarized (n = 95)^23^. Within each group, we compared outcomes between patients treated with carboplatin and paclitaxel alone or in combination with bevacizumab. Consistent with our previous analyses, each 1 SD increase in polarization predicted worse OS under standard chemotherapy (n = 95; HR 1.38, 95% CI 1.02–1.86, p = 0.038; **Table S12**). Importantly, prolonged PFS with bevacizumab was observed only in SecB-polarized tumors (HR 0.36, 95% CI 0.22–0.59, p < 0.001; median PFS 11.1 vs 18.7 months), with no benefit provided to SecA-polarized tumors (HR 0.80, 95% CI 0.48–1.35, p = 0.41; median PFS 17.9 vs 21.6 months, **Figure 7k,l**). The treatment and polarization interaction was significant for PFS (median split HR 0.44, 95% CI 0.22–0.90, likelihood-ratio p = 0.025). In a time-varying analysis, the benefit in SecB tumors was specific to the first 15 months, during which bevacizumab was administered (HR 0.23, 95% CI 0.12–0.44, p < 0.001) but lost thereafter (HR 0.87, p = 0.77; period interaction p = 0.022; **Figure 7m**), whereas SecA tumors showed no period-specific effect (interaction p = 0.19; **Figure 7n**). Together these data suggest that the avascular niches of SecB tumors may make them more reliant on angiogenic signaling, and identify secretory polarization as a candidate biomarker associated with greater bevacizumab benefit, to be tested in prospective cohorts.

## DISCUSSION

Here we show that tumor architecture is a dominant driver of epithelial heterogeneity in HGSC. Secretory epithelial cells span a polarization axis from proliferating, progenitor-like SecA to quiescent, environmentally adapted SecB. This polarization is driven by distance from vasculature and hypoxia rather than by clonal genotype. SecB cells co-organize with reprogrammed macrophages into immune-protected niches primed for dissemination. SecB-enriched tumors have worse outcomes after standard chemotherapy, yet preferentially benefit from bevacizumab, positioning secretory polarization as both a prognostic and a candidate predictive biomarker.

HGSC is characterized by extensive copy number alterations that could produce idiosyncratic, patient-specific phenotypes^25^. Instead, we find secretory polarization broadly reproducible across patient samples: 93.5% of patients with ≥ 100 epithelial cells contain SecA, Intermediate, and SecB cells, all existing within individual clones. The proportion of each state varies by environment, with higher SecB representation in the ascites and in hypoxic microenvironments within solid tissue. Previous findings in glioblastoma have similarly demonstrated that recurrent malignant programs are shaped by hypoxic gradients rather than genotype^22,26^.

However, SecA and SecB do not represent a generic pan-cancer program^27–29^, the previously described therapy-associated stress signature (AP-1/IEG), nor EMT features (SNAI1/2, VIM)^15^. Instead, they recapitulate the secretory lineage of the normal and injured fallopian tube epithelium. SecA cells express the canonical regulatory program of secretory progenitors, including MECOM, SOX17, FBXO21, PBX1, and LGR5^30,31^. SecB cells reflect the keratin and alarmin program of injured mucosal epithelium, marked by KRT7/17/19^32^, LCN2^33^, SLPI^34^, and TACSTD2/TROP2^35^. Consistent with the developmental constraint model of cancer, these findings suggest that the microenvironment selects from states bound to its tissue of origin^36^.

While the secretory polarization axis had not yet been identified, multiple SecB markers have been implicated in HGSC previously. KRT7^24,37,38^, KRT17^39,40^, KRT19^24,41,42^ and its cleaved fragment CYFRA21-1^43,44^, and TROP2^37,45,46^ have been linked to poorer outcomes across HGSC cohorts. Additionally, within HGSC tumors, a spatial program termed differentiated-invasive, marked by elevated LCN2 and TACSTD2, was recently characterized and associated with poorer prognosis^47^. Conversely, SecA-polarized regions likely reflect the spatial domains with high MHC-II expression, elevated antigen presentation (HLA-I and CIITA expression) and increased immune infiltration that have been previously identified as protective in HGSC patients^48–50^. Additionally, secretory polarization may provide a cellular explanation for the link previously made between HGSC molecular subtypes and morphology-based architectural patterns^51^. SecA cells were highly represented in the proliferative and immunoreactive tumors, while SecB cells were abundant in mesenchymal and differentiated subtypes.

SecB-polarized epithelial cells and macrophages assemble into a coherent niche at the luminal interface of papillary growth. Within this hypoxic niche, SecB cells trade proliferation for HIF/NF-κB-driven survival and glycolysis, upregulating MCT-4 (*SLC16A3*), which could acidify the niche and further contribute to lymphocyte exclusion^52,53^. Exclusion was most pronounced for B cells, which are generally associated with tertiary lymphoid structures and with favorable outcomes in HGSC^54^. B cell loss may therefore contribute to the poor prognosis of SecB-enriched tumors. Macrophages were instead retained while adopting a hypoxic, SPP1⁺ MMP9⁺ glycolytic state matching that of the tumor-associated macrophages previously linked to poor outcome across cancers, including HGSC^55^.

SecB cells and hypoxic macrophages appear to engage in reciprocal signaling that sustains the niche. Macrophages supply *VEGFA*, and SecB cells in contact with macrophages further activate NF-κB targets *BIRC3* and *ICAM1*. SecB cells may in turn evade phagocytosis and complement recognition through CD47 and CD55, and may suppress immunity through galectin-9 and IL-32. These cells likely also coordinate adhesion and proteolysis programs within the niche. SecB-adjacent macrophages express cathepsins which can activate local matrix metalloproteinases (MMPs). Notably, CTSL, a marker of SecB adjacent macrophages, can promote matrix remodeling by inhibiting SecB expressed SLPI, which normally functions to protect against injury^56^. SPP1 expressing macrophages can engage CD44 on macrophages and SecB cells and these adhesion molecules in conjunction with elevated ICAM1 likely support multi-cellular anchorage within the niche^57,58^. Together, these programs likely support the establishment of cohesive, multicellular units that detach into ascites^59,60^. As ascites is a hypoxic fluid^61^ and is itself strongly enriched for SecB cells (68% vs 14% in primary tumors), these data support a model in which SecB niches provide a pre-adapted source of disseminating spheroids.

This SecB niche is quiescent and self-sustaining, and therefore may be expected to resist chemotherapy. Indeed, secretory polarization may provide a cellular explanation for why HGSC is often highly sensitive to initial chemotherapy, but recurs almost universally. The high proliferative fraction of SecA cells may render them particularly sensitive to platinum and taxane chemotherapy, contributing to this initial response. Conversely, SecB cells are predominantly in G1 and reside in poorly perfused regions that may limit drug delivery, which could favor persistence. Consistent with this, SecB proportions increased in post-chemotherapy primary tumors and rose after treatment in 12 of 18 patient-matched samples across various tumor sites. SecB-enriched tumors also had shorter survival across three independent cohorts. SecB cells may therefore represent a reservoir of drug-tolerant persisters that seed recurrence, a hypothesis testable by longitudinal sampling.

Secretory polarization may also explain the longstanding asymmetry in HGSC response to bevacizumab, which improves PFS in unselected patients but rarely OS. In ICON7^23^, the PFS benefit of bevacizumab was confined to SecB-polarized tumors (HR 0.36; median 11.1 vs 18.7 months), with no significant benefit in SecA-polarized tumors, and was confined to the first 15 months, spanning treatment. SecB niches are poorly perfused and hypoxic yet enriched with macrophage-derived VEGFA, suggesting that they depend on ongoing angiogenic signaling. Anti-VEGF therapy may therefore suppress SecB niches only while administered, allowing them to re-emerge after withdrawal and accounting for PFS gains that do not translate into OS. Unlike prior candidate biomarkers for bevacizumab response, secretory polarization proposes a multi-cellular mechanism linked to clinical response, and therefore the degree of polarization could guide patient selection, maintenance duration, and inform combination therapies.

The fact that this state was not characterized previously can likely be attributed to the experimental models currently used to study HGSC. SecB was largely lost in our PDOs, which collapsed toward SecA, mirroring the previously reported collapse of basal and classical pancreatic cancer organoids into a default classical-like state^21^. Likewise, neither published HGSC organoid or spheroid models, including those cultured under hypoxic conditions, nor the KPCA.B syngeneic model could recapitulate both the hypoxic and quiescent features of SecB. Furthermore, no environmental signal or culturing media composition tested was able to polarize cells to SecB within our PDOs, and only IFN-γ induced any meaningful SecB gene expression. This partial success with IFN-γ suggests that immune-derived signals may play a role in initiating polarization early in niche formation, prior to the establishment of the lymphocyte-excluding acidic and hypoxic cores. Altogether, these results suggest current preclinical platforms may be systematically biased toward the proliferative, chemo-sensitive SecA state, and screening these would underrepresent the population most likely to persist after chemotherapy. New models incorporating hypoxic gradients, immune signals and multicellular organization will be required to effectively study and target the SecB niche.

While we currently lack the complex systems required to model SecB, secretory polarization is already measurable with tools used in pathology. For example, KRT7 and KRT19 protein abundance tracked Xenium-defined polarization in the same TMA cores, whereas non-SecB epithelial markers (E-cadherin, KRT18, VIM) did not. This is consistent with prior reports that KRT7 and KRT19 independently predict worse survival^24^. Beyond tissue, serum CYFRA 21-1, a cleaved KRT19 fragment, may offer a minimally invasive readout of SecB burden^43^. There are therefore multiple potential strategies for testing polarization-based stratification prospectively without single-cell or spatial profiling. Prospective validation of these strategies against chemotherapy and bevacizumab outcomes will be required to translate these hypotheses to clinical application.

This study has several limitations that warrant future validation. The spatial analyses are observational and cannot establish whether tissue architecture induces, selects, or maintains secretory states. Hypoxia, metabolic activity, and signaling were inferred primarily from transcriptional expression rather than measured directly. Finally, clinical associations were examined retrospectively and the utility for treatment selection requires formal prospective validation.

Together, our findings position tissue organization as a dominant driver of epithelial heterogeneity in HGSC, shaping malignant cell states constrained by the developmental history of the cancer. These malignant states reciprocally interact with the TME, forming heterogeneous niches that emerge as recurrent units throughout the tumor and may drive distinct aspects of disease biology. Current cytotoxic therapy preferentially targets the proliferative SecA compartment, whereas the transient benefit of bevacizumab in SecB-polarized tumors suggests that the SecB niche is targetable by anti-VEGF therapy. Durable improvement of HGSC outcomes will likely require biomarker-guided use of existing agents combined with approaches that directly engage the SecB niche. Finally, disrupting the signaling that sustains the niche, as bevacizumab likely does, may prevent its persistence altogether.

## Acknowledgements

We thank the molecular pathology core facility of the CRCHUM.

The authors would like to acknowledge the assistance of StemCore Laboratories Genomics Core Facility (OHRI, uOttawa), RRID:SCR_012601.

## SUPPLEMENTARY FIGURE LEGENDS

**Figure S1.**
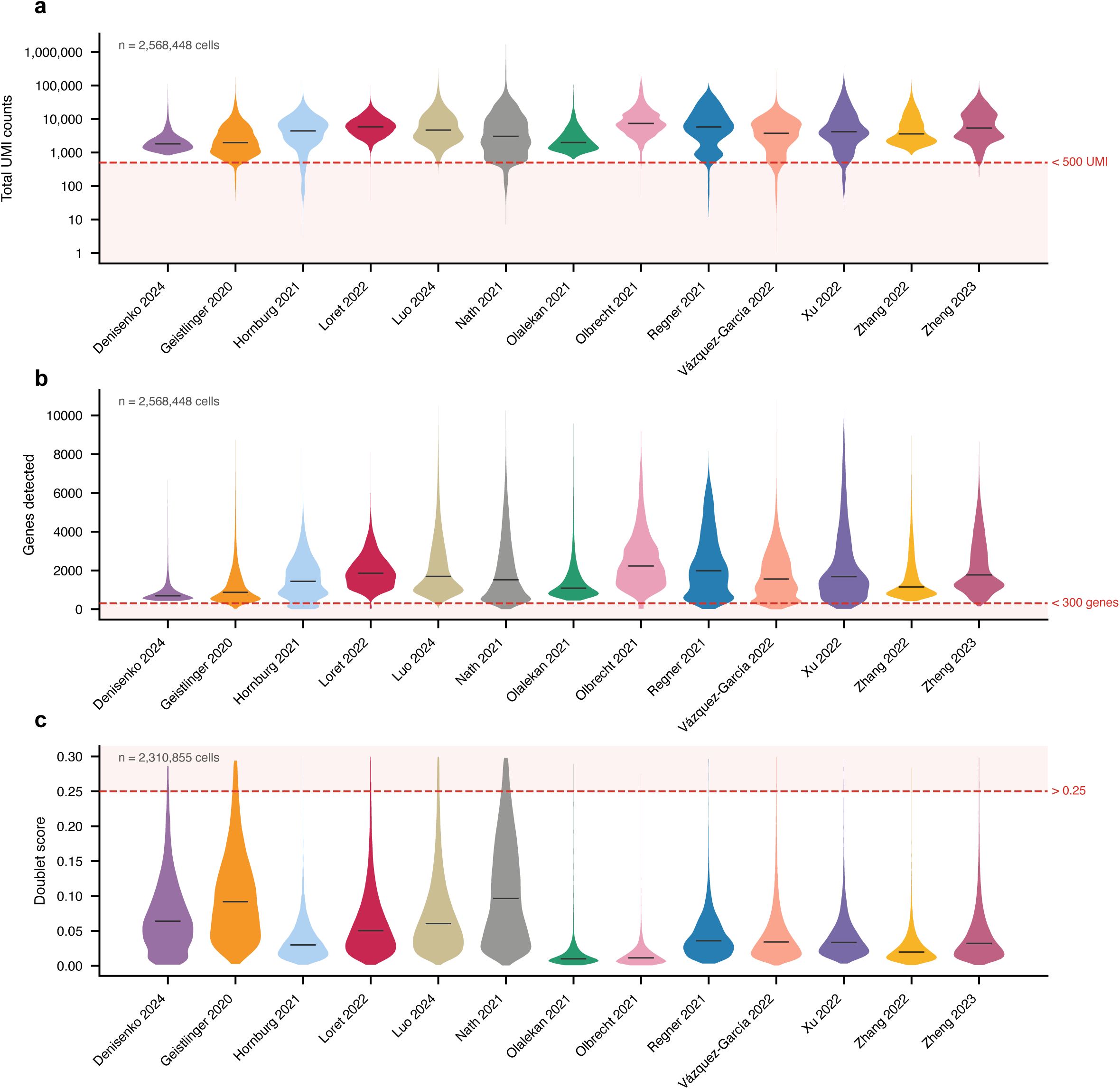
Quality control metrics by study across our integrated HGSC scRNA-seq atlas. **(a**) Violin plots of total UMI counts (log scale) per cell by study; red dashed line demonstrates the threshold applied across studies (<500 UMIs). (**b**) Violin plots of genes detected per cell by study; red dashed demonstrates the threshold applied (<300 genes). (**c**) Violin plots of Scrublet doublet scores per cell by study; red dashed line demonstrates the threshold applied across studies (> 0.25). Horizontal bars represent the study medians.

**Figure S2.**
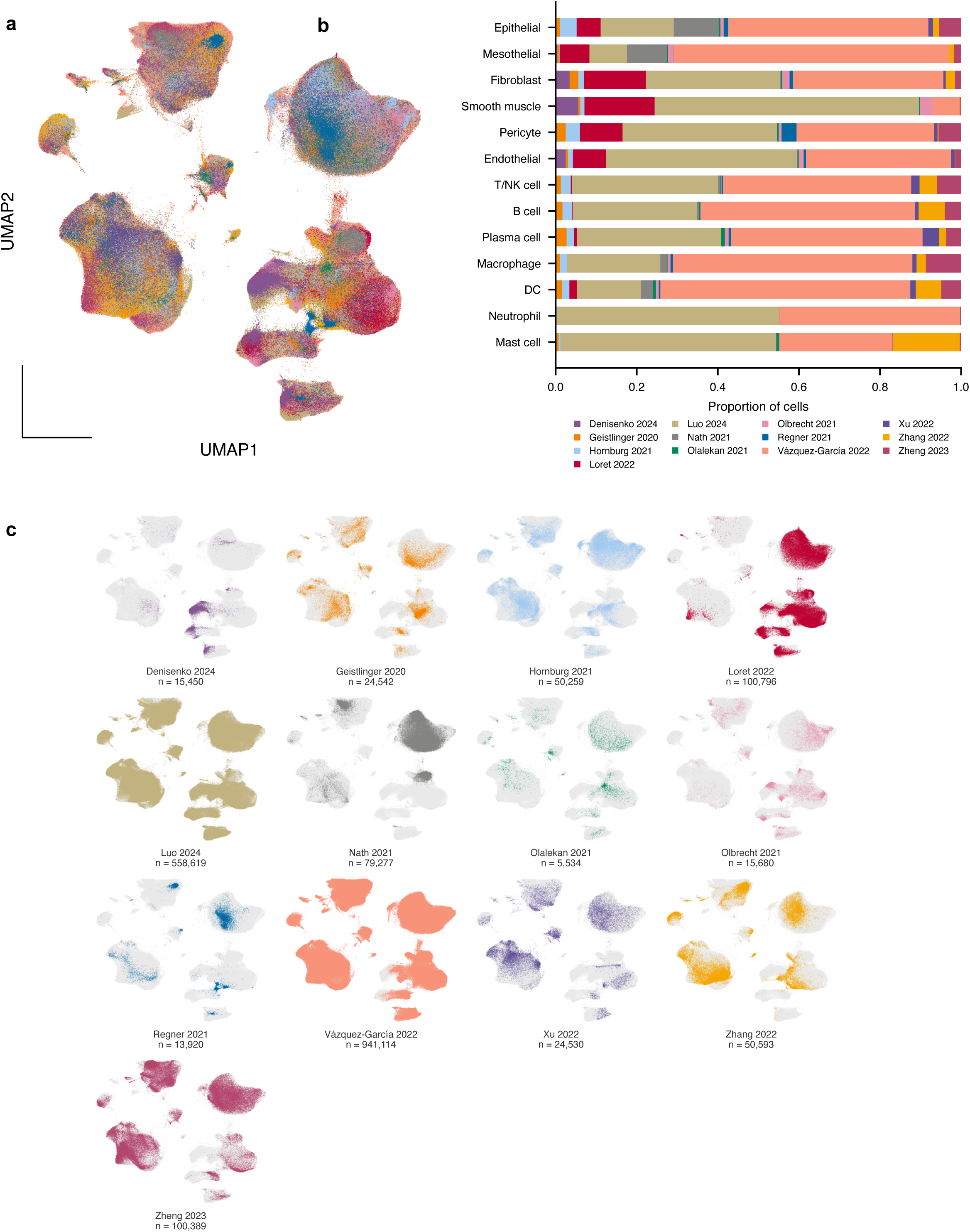
Contribution of each study to our integrated HGSC scRNA-seq atlas. (**a**) UMAP of our integrated scRNA-seq HGSC atlas (n = 1,980,703 cells), colored by study. (**b**) Proportional stacked bar graph demonstrating the contribution of each study to each of the level 1 cell types (n = 13). (**c**) UMAP visualizations highlighting the cells contributed, colored by study, within the integrated HGSC scRNA-seq atlas.

**Figure S3.**
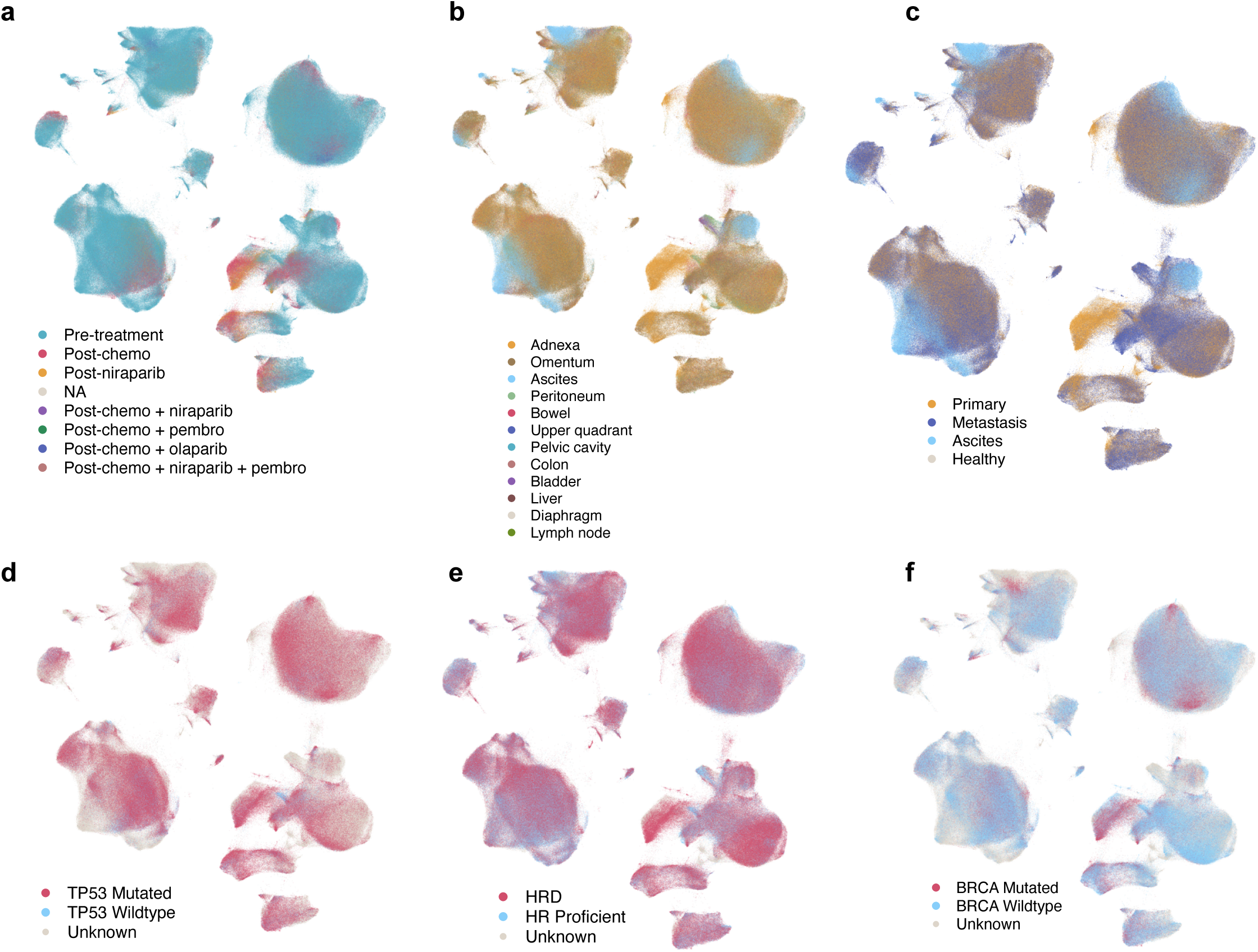
Clinical and genomic metadata across the integrated HGSC scRNA-seq atlas. UMAP of the integrated HGSC scRNA-seq atlas colored by (**a**) treatment status, (**b**) anatomic site, (**c**) broad metastatic site, (**d**) TP53 status, (**e**) HRD status, (**f**) BRCA status.

**Figure S4.**
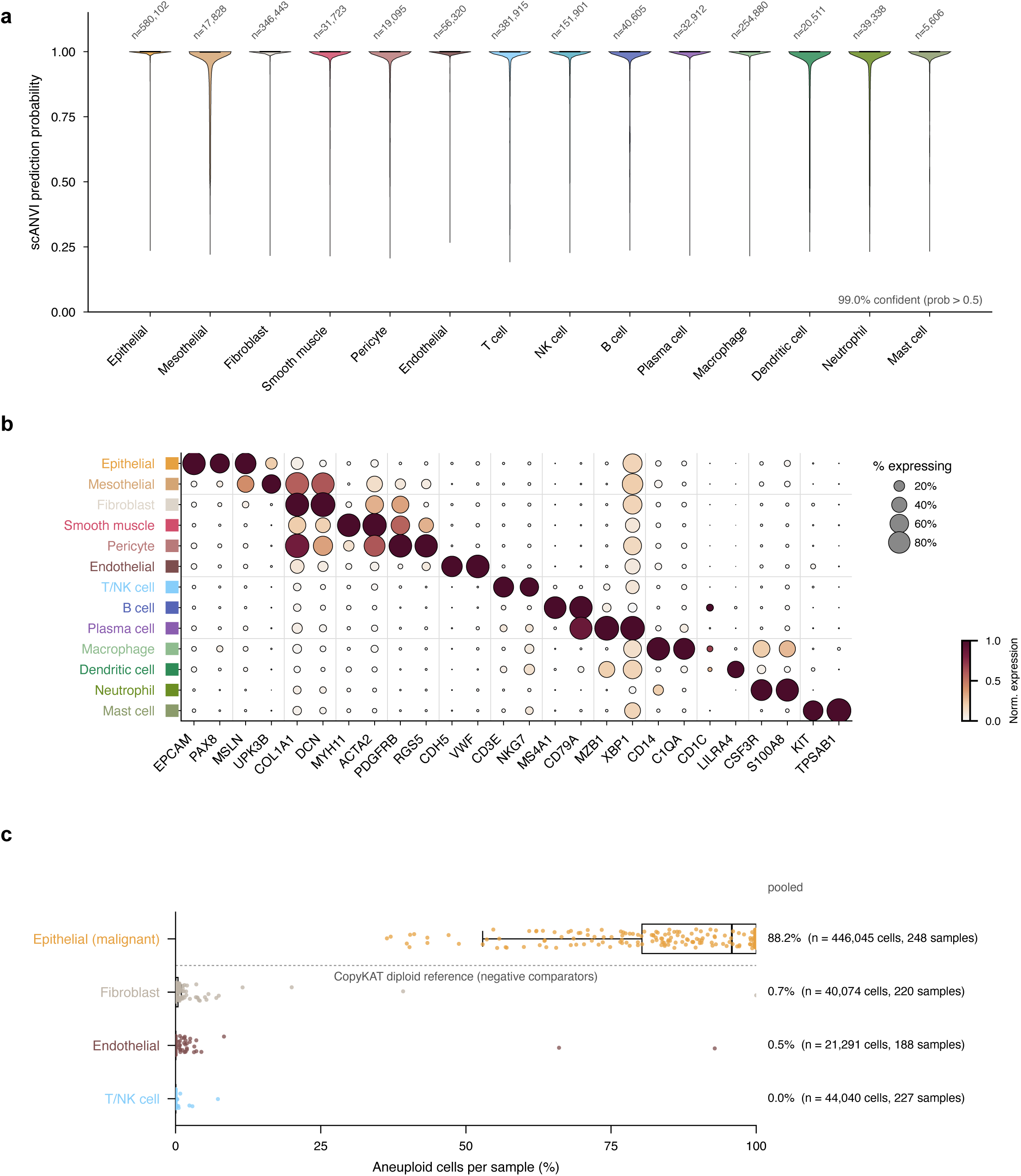
Cell type annotation validation within the HGSC scRNA-seq atlas. (**a**) Violin plots of scANVI prediction probability for each scANVI-predicted cell class (14 cell types, T and NK cell shown separately) with the number of cells per cell type. (**b**) Dot plot of canonical cell type markers across the 13 annotated cell types, with color representing the scaled mean expression and size representing the percentage of expressing cells. (**c**) Box plot demonstrating the percentage of CopyKAT-inferred aneuploid cells per sample across the epithelial, fibroblast, endothelial and T/NK cell compartments. Points represent all samples with a minimum of 20 cells in each group and boxes represent the median and interquartile range; pooled percentages are quantified and shown in the right.

**Figure S5.**
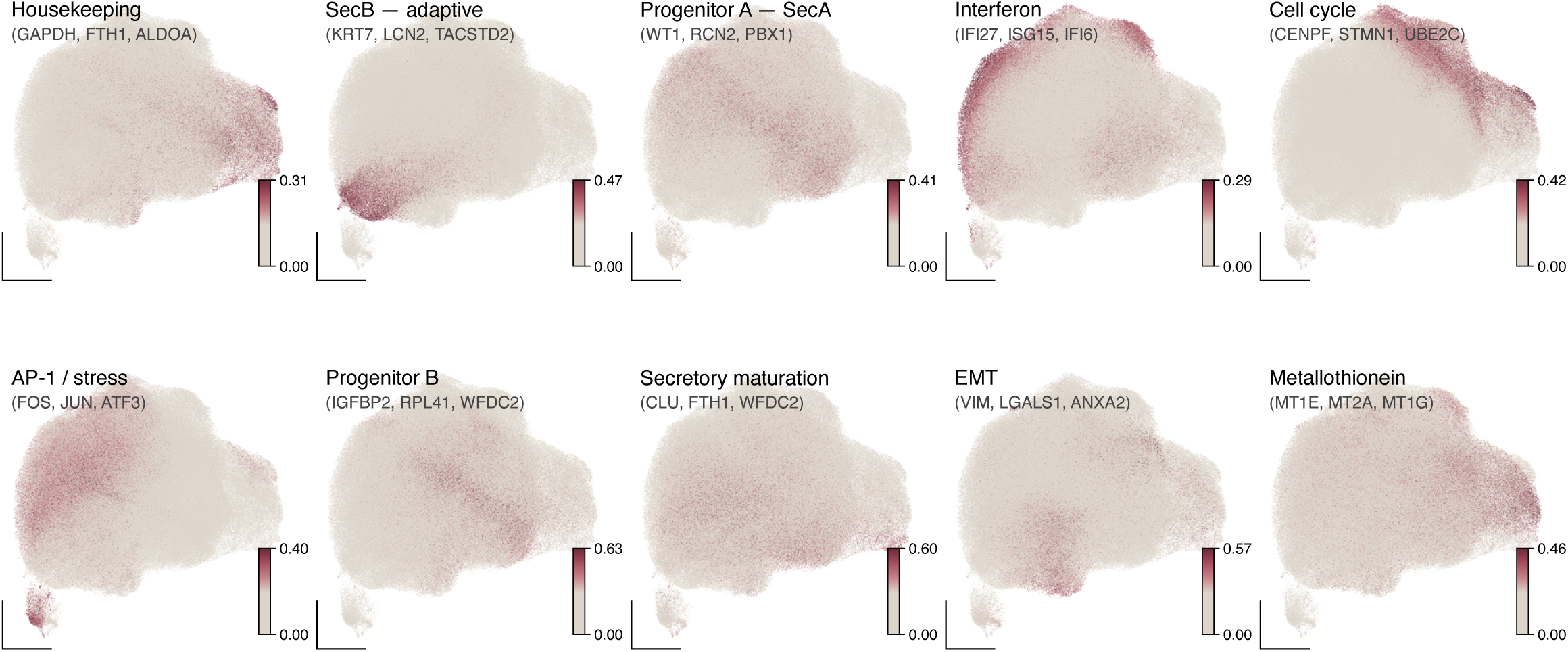
NMF factor usage across the integrated epithelial compartment. UMAP of the integrated epithelial compartment (n = 575,366) with per-cell usage of each of the 10 NMF factors visualized by factor usage. Factors are labelled by inferred program and top marker genes.

**Figure S6.**
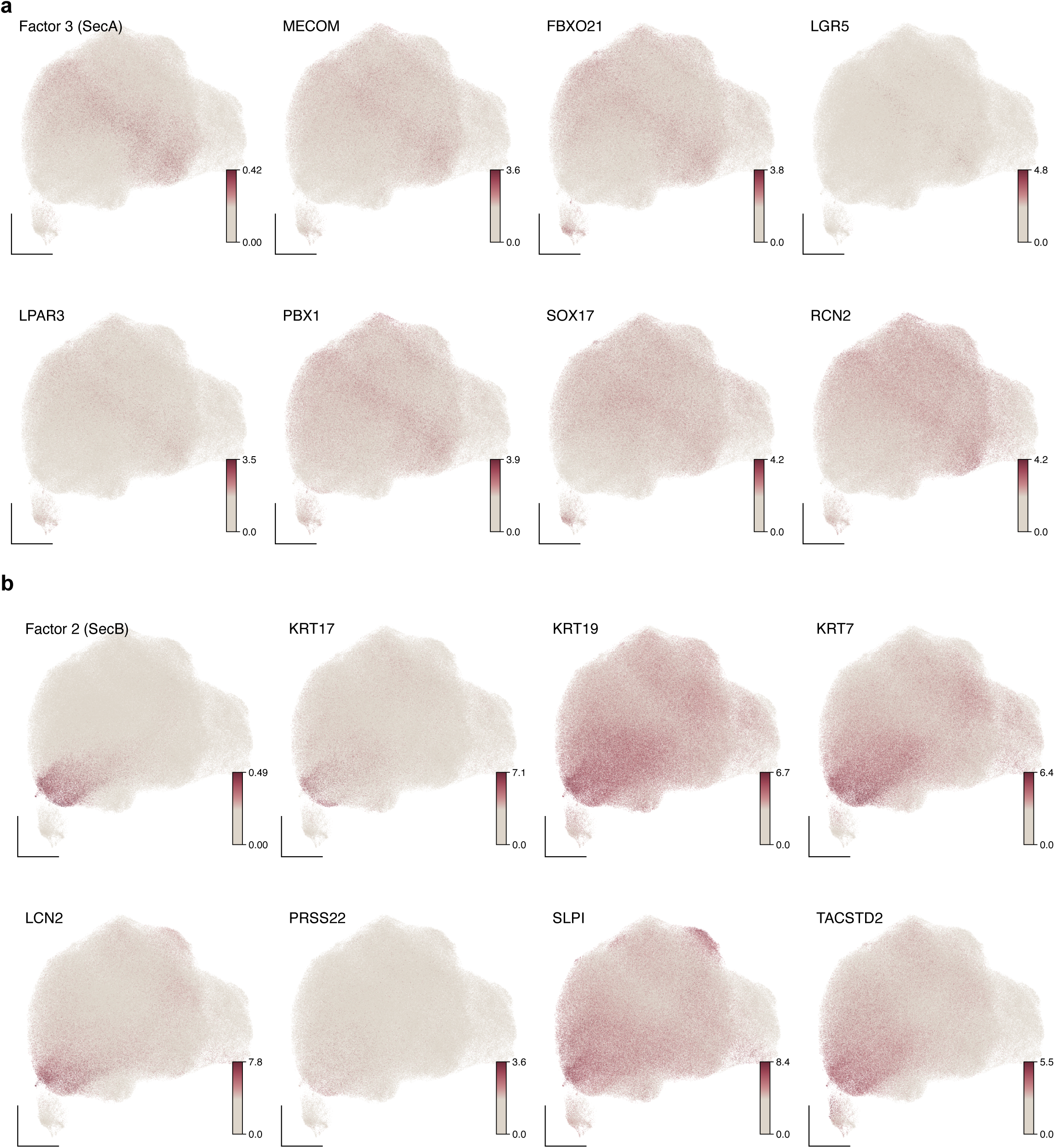
SecA and SecB signature gene expression across the integrated epithelial compartment. UMAP of the integrated epithelial compartment (n = 575,366) visualizing (**a**) SecA NMF factor loading and SecA marker genes (MECOM, SOX17, FBXO21, PBX1, LGR5, LPAR3, RCN2). (**b**) SecB NMF factor loading and SecB marker genes (KRT17, LCN2, TACSTD2, KRT7, KRT19, SLPI, PRSS22).

**Figure S7.**
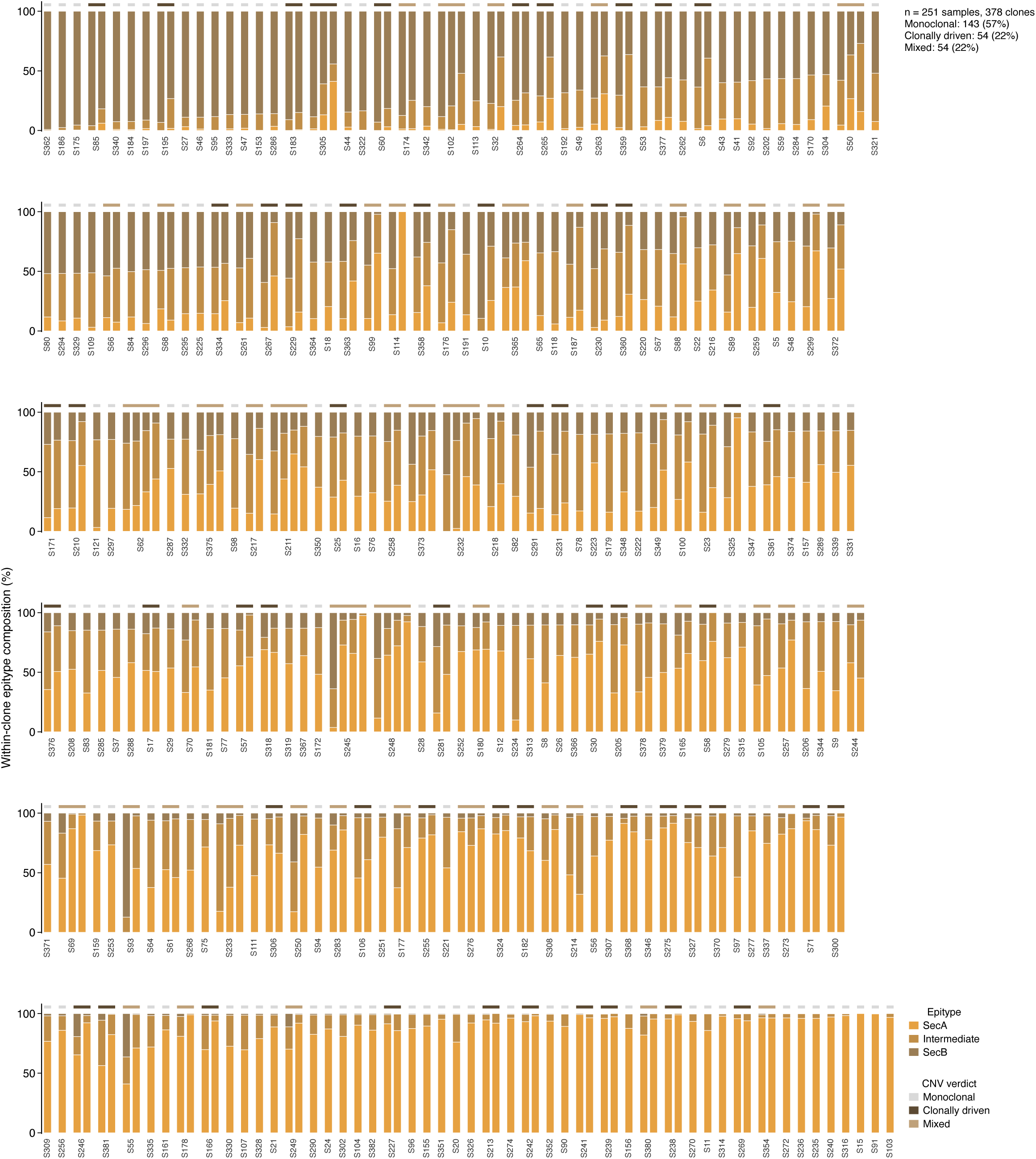
SecA, Intermediate and SecB epithelium coexist within CopyKAT-inferred clones. Stacked bar graph demonstrating the proportion of SecA, Intermediate and SecB within each CopyKAT-inferred subclone with a minimum of 20 cells (n = 378 subclones across 251 samples; 143 monoclonal, 54 clonally driven and 54 mixed), grouped by sample and ordered by sample-level SecB proportion. The banner above represents the clonality for the sample (gray = monoclonal, brown = polyclonal, beige = mixed).

**Figure S8.**
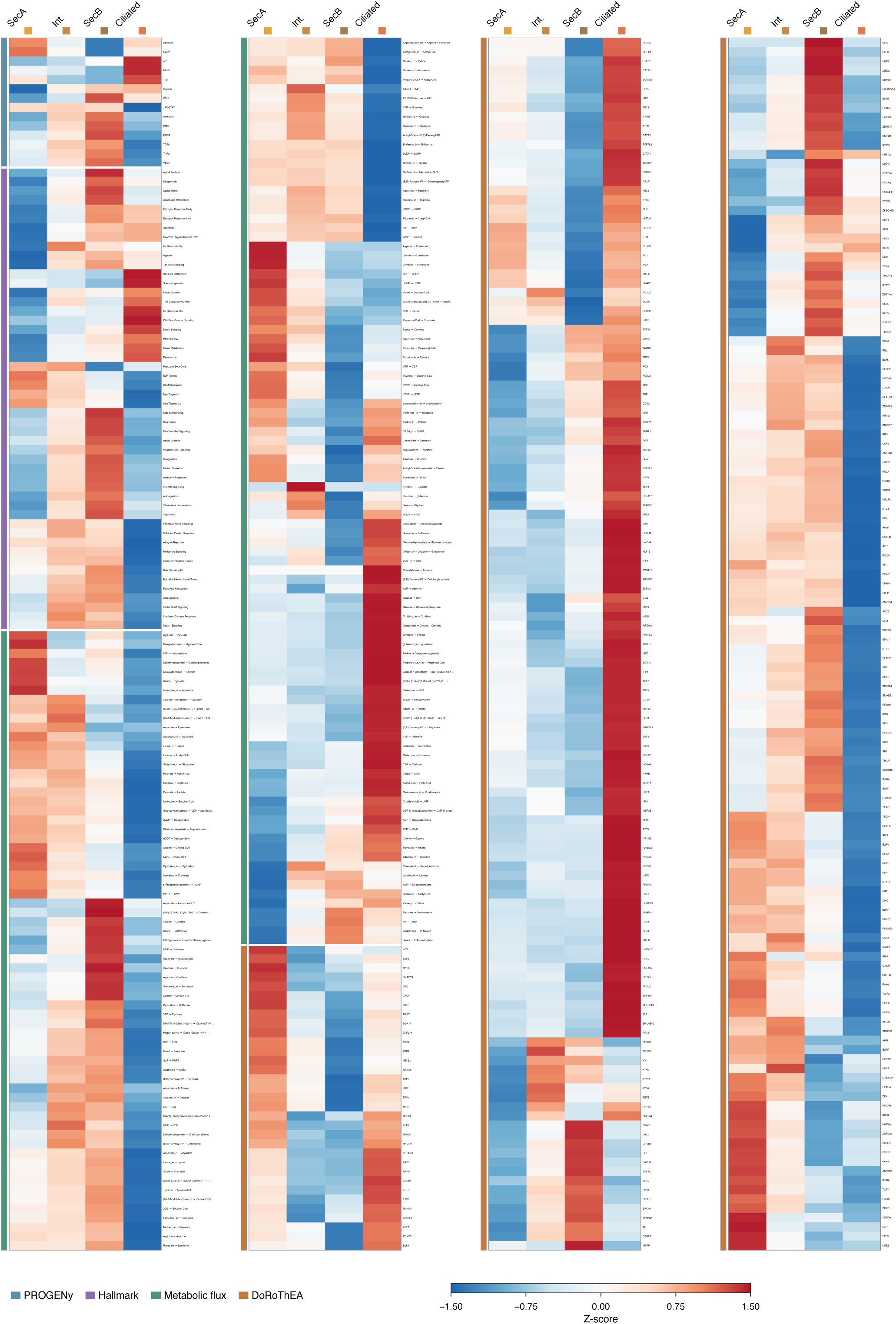
Pathway, metabolic, and transcription factor activity across epithelial states. Heatmaps of row-normalized z-scores for SecA, Intermediate, SecB and ciliated epithelium of inferred pathway activity (PROGENy, 14 pathways), gene set enrichment (Hallmark, 50 gene sets), metabolic flux estimation (scFEA, 165 modules), and transcription factor activity (DoRothEA, 294 regulons).

**Figure S9.**
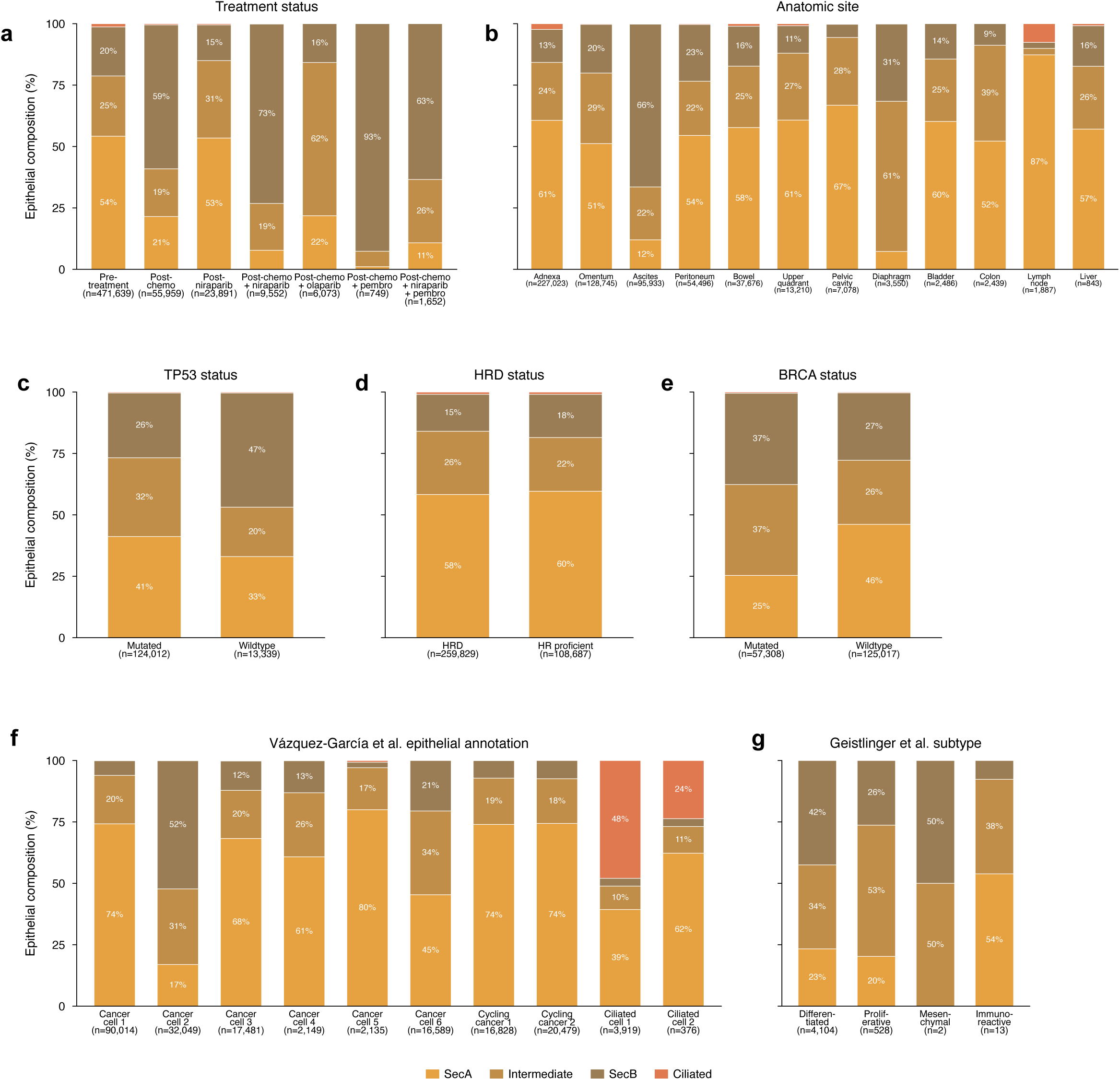
Composition of the epithelial compartment states across clinical, genomic, and previously annotated epithelial clusters. Stacked bar graphs demonstrating the proportion of SecA, Intermediate, SecB and ciliated epithelium by (**a**) treatment status (**b**) anatomic site, (**c**) TP53 status, (**d**) HRD status, (**e**) BRCA status, (**f**) Vázquez-García et al. epithelial cluster annotation (SPECTRUM cohort) and (**g**) Geistlinger et al. consensusOV molecular subtype. The number of epithelial cells is annotated, and percentages are shown for those greater than 8%.

**Figure S10.**
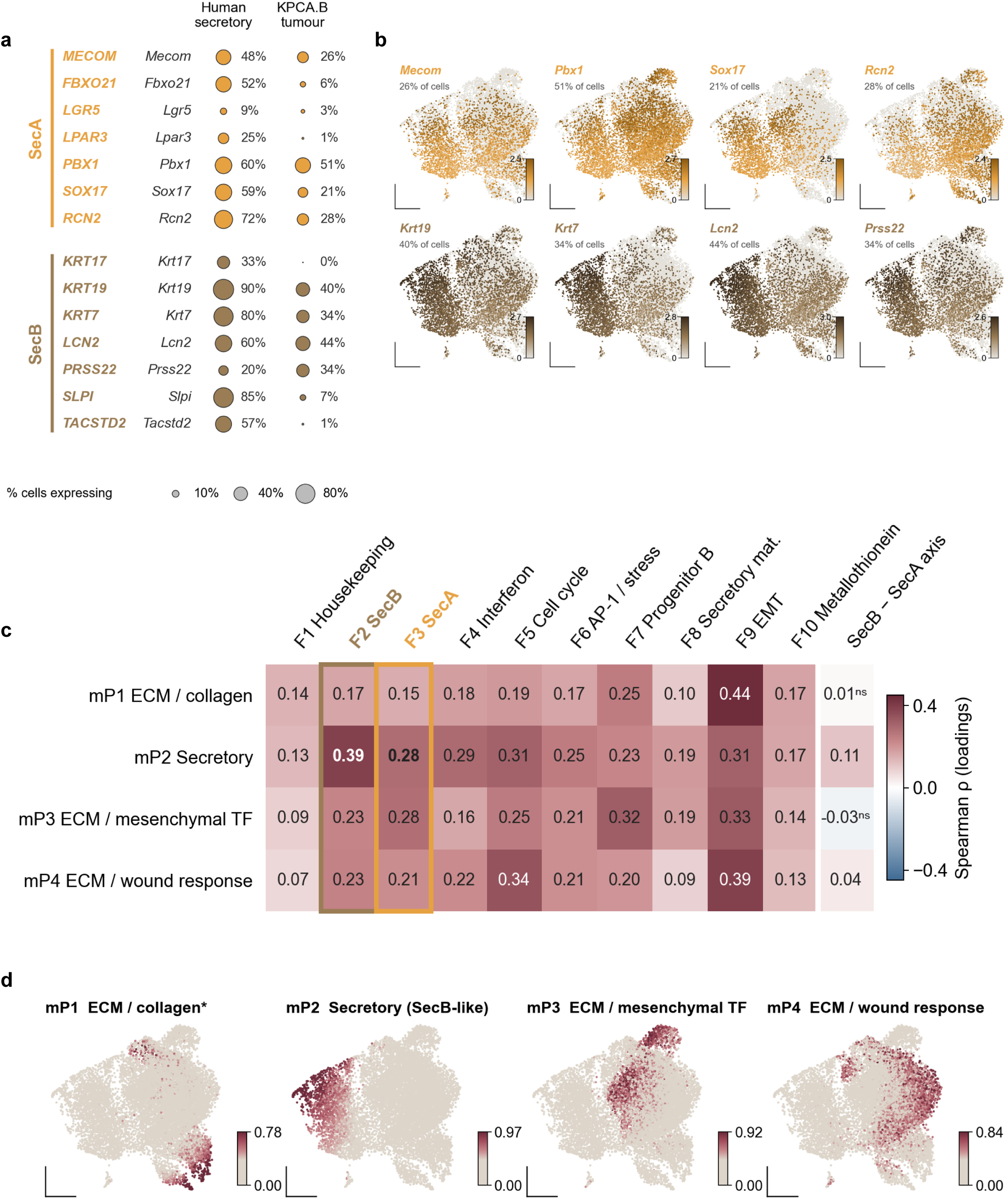
Programs of KPCA.B mouse tumor epithelium compared with human SecA and SecB programs. (**a**) Dot plot demonstrating the SecA and SecB markers and how each translates from the HGSC human scRNA-seq atlas to the mouse orthologs within the KPCA.B tumor epithelium (n = 12,429 cells). (**b**) UMAP of KPCA.B tumor epithelium visualizing expression of SecA and SecB markers expressed in more than 20% of cells. (**c**) Heatmap demonstrating Spearman correlation values between the gene loadings of the four NMF programs of KPCA.B tumor epithelium compared to the human NMF factors and the human SecB and SecA polarizations score, ns represents FDR > 0.05 (gene-based permutation). (**d**) UMAP of KPCA.B tumor visualizing the per-cell usage of each NMF factors.

**Figure S11.**
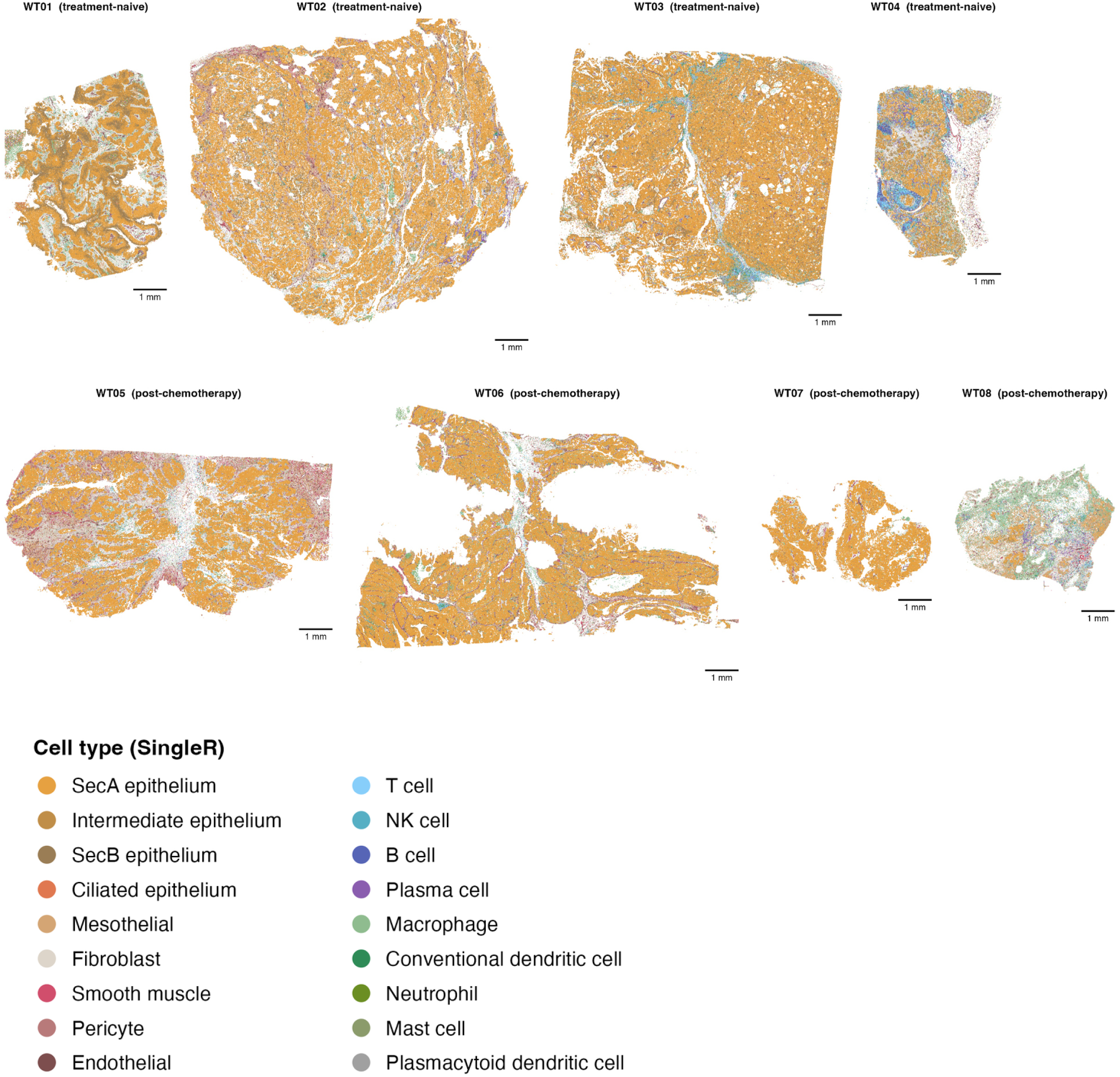
Cell type annotation across our Xenium whole-tissue cohort. Whole-tissue spatial maps of the eight HGSC tissue sections (WT01 – WT04 treatment naïve, WT05 – WT-08, post-chemotherapy, n = 1,898,772 cells) colored by SingleR cell type annotation (18 cell types). Scale bar represents 1 mm.

**Figure S12.**
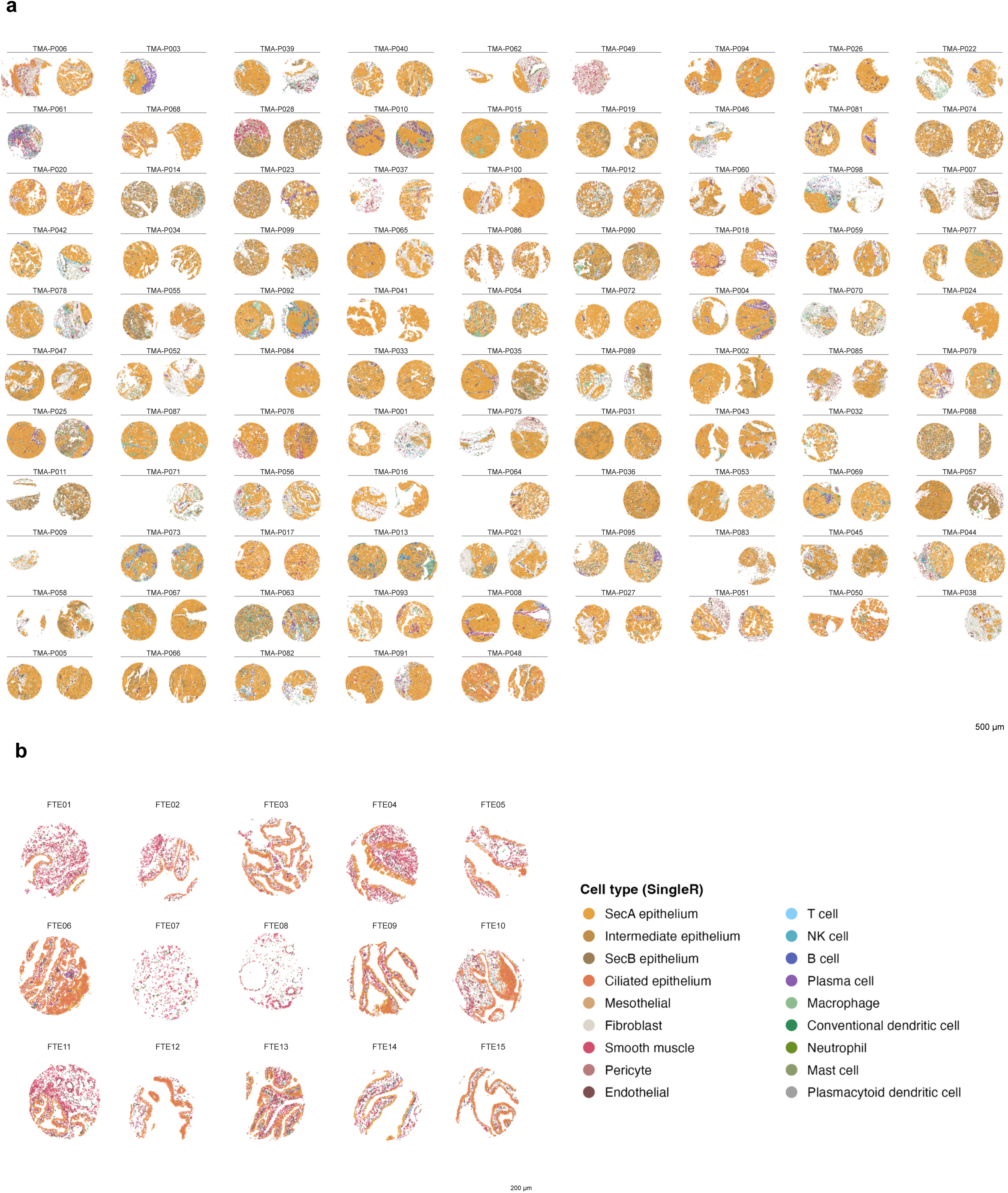
Cell type annotation across our Xenium TMA A and fallopian tube epithelium cores. Spatial maps of (**a**) TMA A (95 patients, 177 cores) and (**b**) fallopian tube epithelium cores (n = 15 cores, n = 34,281 cells), colored by SingleR cell type annotation (18 cell types). Scale bars represent (**a**) 500 µm and (**b**) 200 µm.

**Figure S13.**
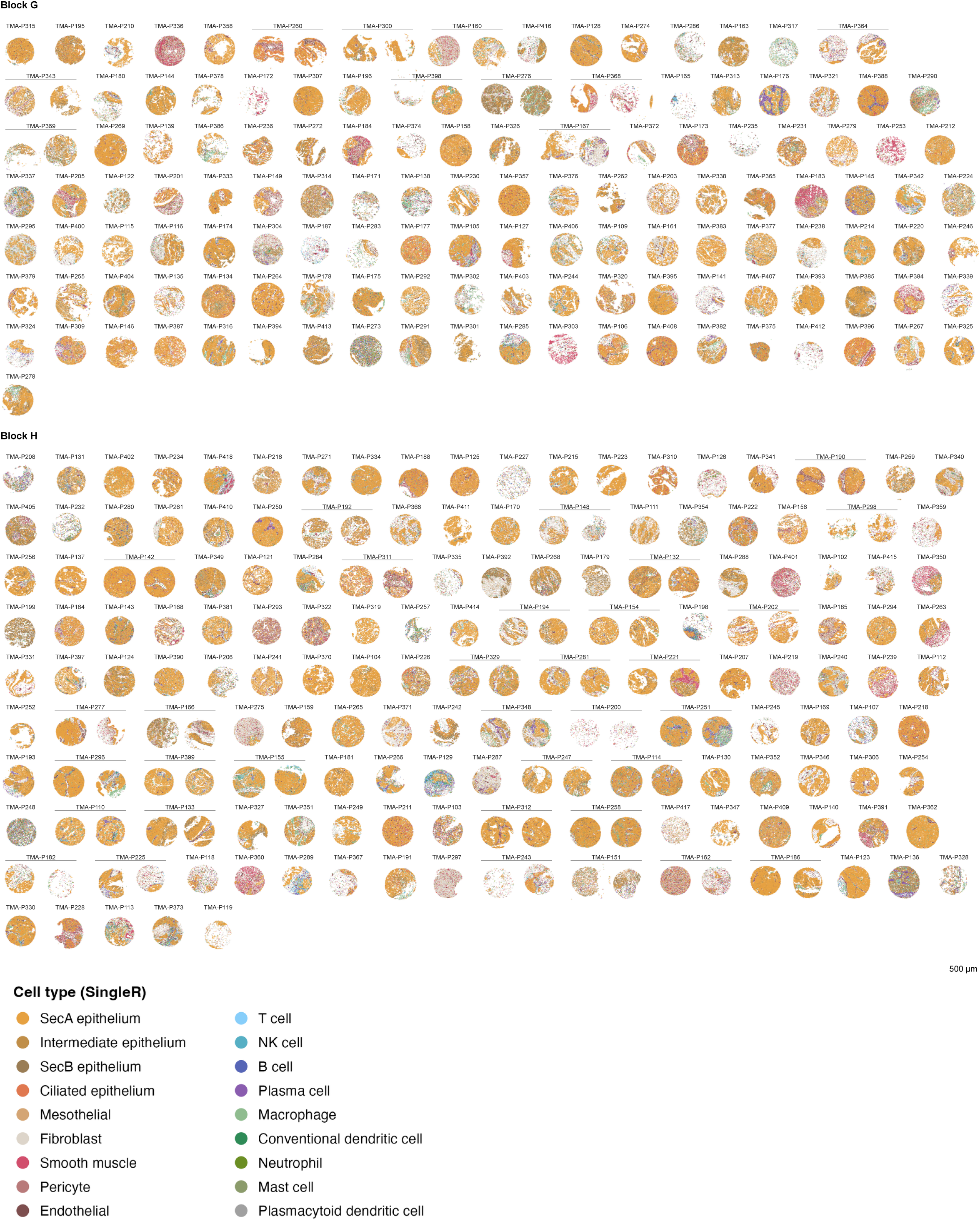
Cell type annotation across our Xenium TMA. **B.** Spatial maps of TMA B (283 patients, 326 cores with 43 two-core pairs) colored by SingleR cell type annotation (18 cell types). Scale bar represents 500 µm.

**Figure S14.**
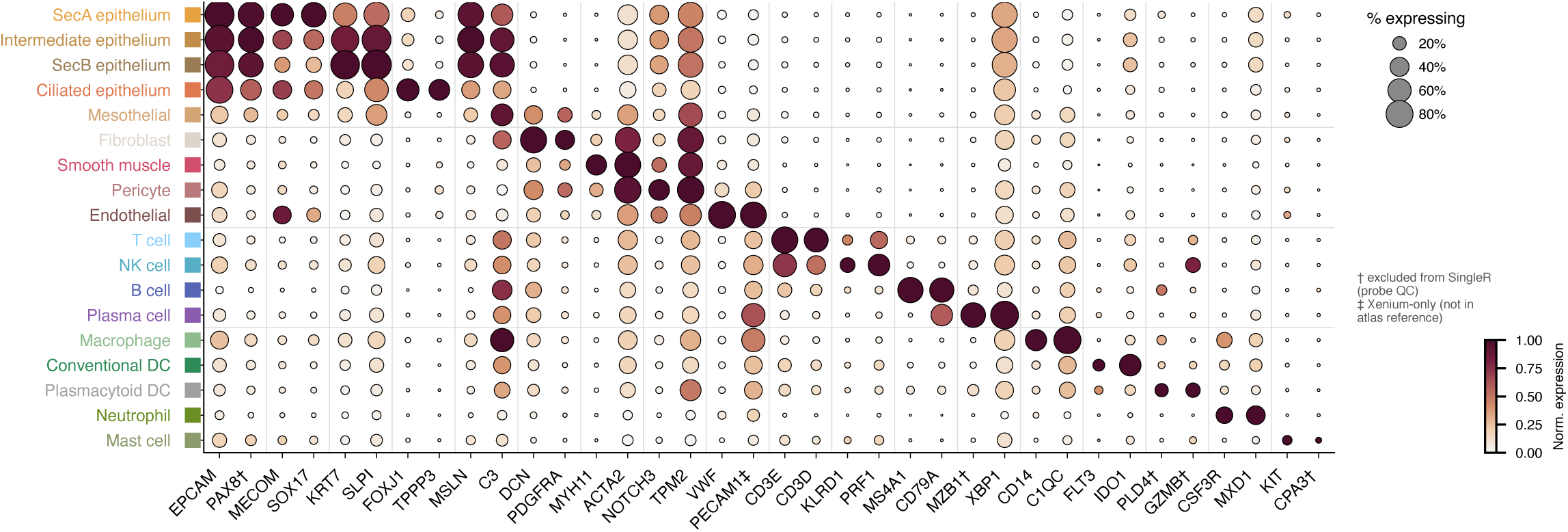
Cell type annotation validation across our Xenium cohort. Dot plot of canonical cell type markers across the 18 annotated cell types, with color representing the scaled mean log-normalized expression and size representing the percentage of expressing cells, † denotes genes excluded from SingleR by probe QC and ‡ denotes a Xenium-only gene absent from the atlas reference.

**Figure S15.**
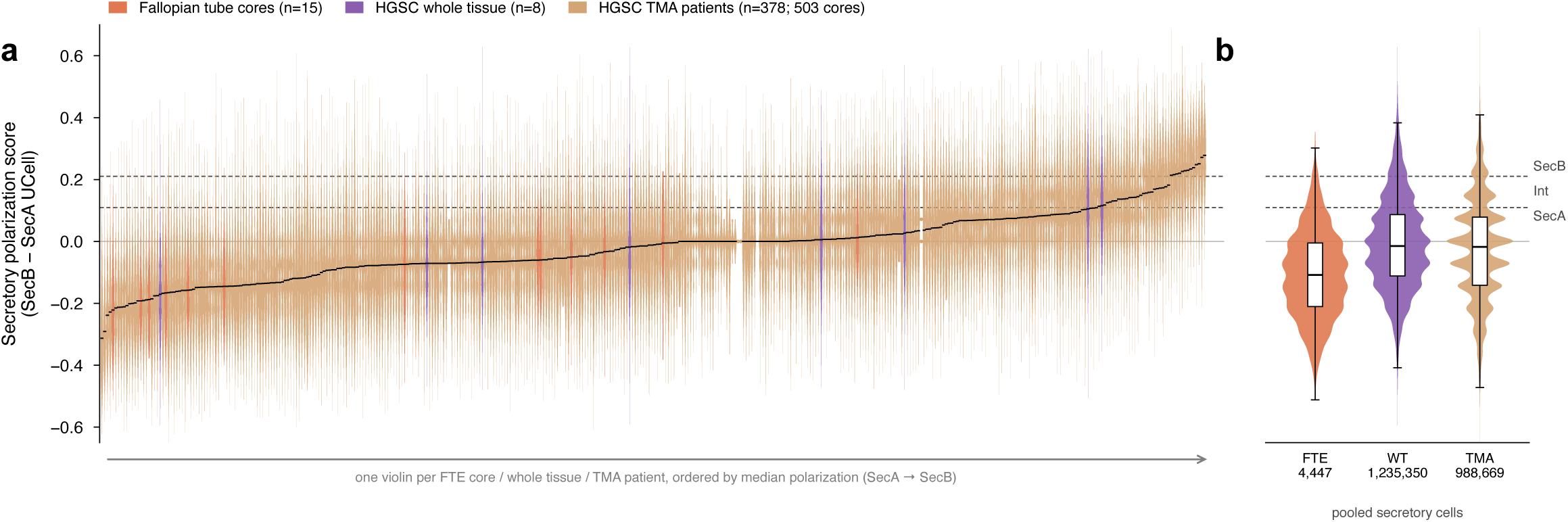
Distribution of secretory polarization across the Xenium cohort. (**a**) Violin plots of the secretory polarization score (SecB – SecA UCell score) for fallopian tube epithelium cores (n = 15, orange), HGSC whole tissue (n = 8, purple) or across our TMAs (n = 378 patients, 503 cores). (**b**) Pooled distributions of all secretory cells per cohort. Dashed lines represent the polarization thresholds separating SecA, Intermediate and SecB cells. HGSC units versus FTE cores: two-sided Mann–Whitney test on per-unit median polarization

**Figure S16.**
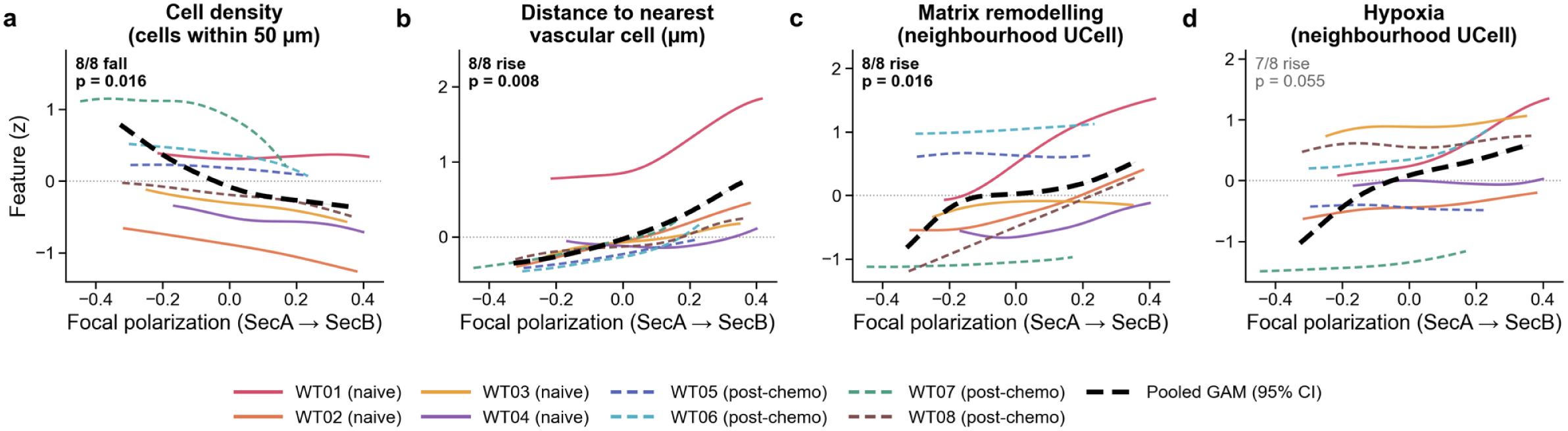
Per-tissue validation of environmental features along the secretory polarization axis. Generalized additive models of (**a**) cell density, (**b**) distance to nearest vascular cell (µm), (**c**) Matrix remodeling score (UCell) and (**d**) neighborhood hypoxia score (UCell). Colored lines represent individual whole-tissue fits on all cells of each tissue (n = 8; WT01–WT04 treatment-naïve, solid; WT05–WT08 post-chemotherapy, dashed). The black dashed line represents the pooled model across all 1,235,350 secretory cells with each tissue weighted equally (95% CI), which reflects both within- and between-tissue variation. Each panel reports the number of tissues changing in the pooled direction and the p value of a two-sided paired Wilcoxon signed-rank test comparing the top and bottom 20% of polarization within each tissue. Curves are shown over the 1st–99th percentile of polarization.

**Figure S17.**
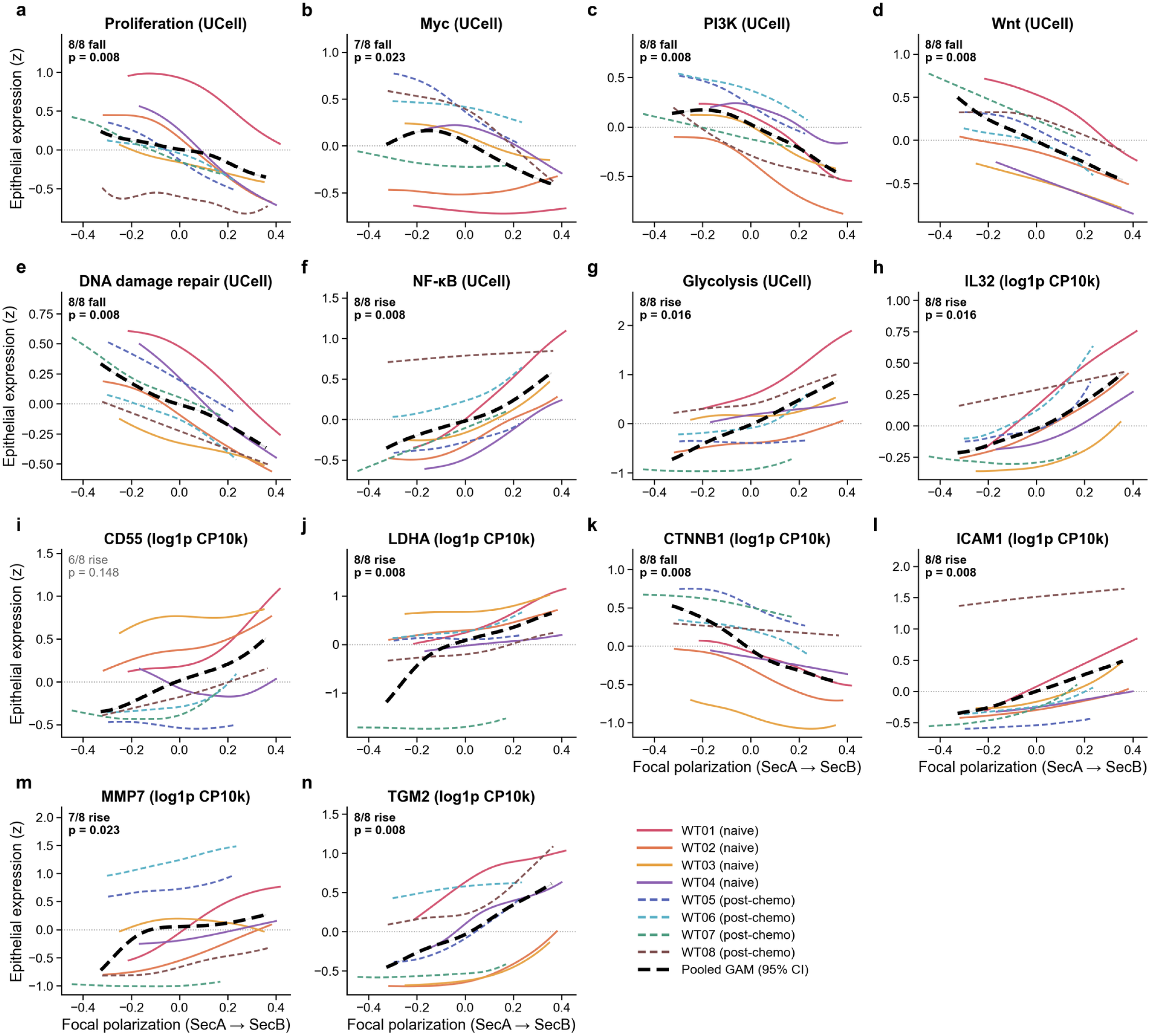
Per-tissue validation of epithelial-intrinsic expression along the secretory polarization axis. Generalized additive models of epithelial UCell scores (z-scored) for (**a**) proliferation, (**b**) Myc, (**c**) PI3K, (**d**) Wnt, (**e**) DNA damage repair (BRCA1, BRCA2, ATM, CHEK2), (**f**) NF-κB and (**g**) glycolysis, and expression (log1p CP10k, z-scored) of (**h**) *IL32*, (**i**) *CD55*, (**j**) *LDHA*, (**k**) *CTNNB1*, (**l**) *ICAM1*, (**m**) *MMP7* and (**n**) *TGM2* in secretory epithelial cells as a function of focal cell polarization score. Colored lines represent individual whole-tissue fits on all cells of each tissue (n = 8; WT01–WT04 treatment-naïve, solid; WT05–WT08 post-chemotherapy, dashed). The black dashed line represents the pooled model across all 1,235,350 secretory cells with each tissue weighted equally (95% CI), which reflects both within- and between-tissue variation. Each panel reports the number of tissues changing in the pooled direction and the p value of a two-sided paired Wilcoxon signed-rank test comparing the top and bottom 20% of polarization within each tissue. Curves are shown over the 1st–99th percentile of polarization.

**Figure S18.**
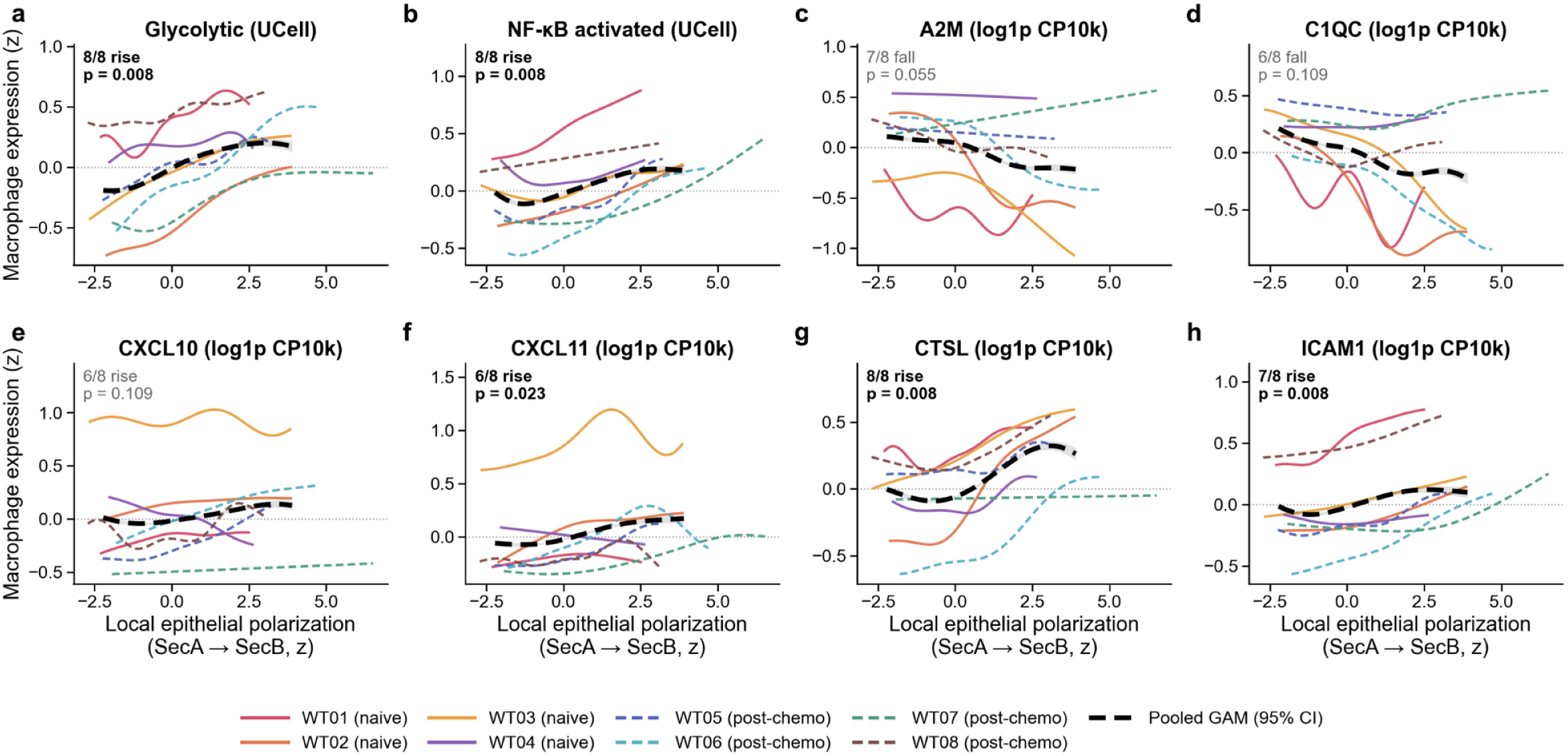
Per-tissue validation of macrophage-intrinsic expression along the secretory polarization axis. Generalized additive models of macrophage UCell program scores (z-scored) for (**a**) glycolysis and (**b**) NF-κB activation, and expression (log1p CP10k, z-scored) of (**c**) *A2M*, (**d**) *C1QC*, (**e**) *CXCL10*, (**f**) *CXCL11*, (**g**) *CTSL* and (**h**) *ICAM1* in macrophages as a function of local epithelial polarization within 50 µm. Colored lines represent individual whole-tissue fits on all cells of each tissue (n = 8; WT01– WT04 treatment-naïve, solid; WT05–WT08 post-chemotherapy, dashed). The black dashed line represents the pooled model across all macrophages. Each panel reports the number of tissues changing in the pooled direction and the p value of a two-sided paired Wilcoxon signed-rank test comparing the top and bottom 20% of polarization within each tissue. Curves are shown over the 1st–99th percentile of polarization.

## METHODS

### RESOURCE AVAILABILITY

Lead contact. David Cook

Materials availability. No unique reagents were generated.

Data and code availability. All scRNA-seq datasets used in atlas construction are publicly available. Processed atlas objects, Xenium, Visium HD, PDO snRNA-seq and KPCA.B scRNA-seq have been deposited at [TBD]. The code to reproduce analyses and data visualization is available through https://www.github.com/cook-lab/hgsc-malignant-states.

### EXPERIMENTAL MODEL AND STUDY PARTICIPANT DETAILS

Human Tissues. De-identified archival tissues were used under REBs #20240175-01H and #20240096-01H, with informed consent. We profiled three sources of tissues: eight FFPE HGSC whole tissues (WT01 – WT08), TMA A (101 patients, CHUM cohort) and TMA B (317 patients profiled, COEUR cohort) (**Table S8**).

Patient-derived organoids (PDOs). Eight HGSC organoids were used: OPTO98, OPTO112, OPTO129, and PDO66 from primary tumors and OCAD93, OCAD96, OCAD97 and OCAD106 from ascites. These were obtained from Princess Margaret Cancer Centre and Western University. Organoids were embedded in Matrigel (20,000– 50,000 cells per 50 µL dome). The culture medium was Advanced DMEM/F-12 containing GlutaMAX (2 mM), HEPES (10 mM), antibiotic-antimycotic (100 U/mL), B27 (1×), N-acetyl-L-cysteine (1.25 mM), EGF (20 ng/mL), bFGF (100 ng/mL), FGF-10 (100 ng/mL), Noggin (100 ng/mL), nicotinamide (1 mM), Y-27632 (10 µM), forskolin (10 µM) and β-estradiol (200 nM). The PDO66 medium was further supplemented with N2 (1×), SB431542 (0.5 µM) and BMP2 (10 ng/mL). Organoids were passaged every 1–4 weeks after TrypLE dissociation.

Mice. KPCA.B cells were kindly provided by Dr. Robert Weinberg^62^ and 150,000 cells were implanted orthotopically into the ovarian bursa of C57BL/6 mice under protocol OHRIe-3632. Tumors were collected for scRNA-seq at day 34.

### METHODS DETAILS

#### Single-cell atlas construction

We integrated raw counts from 13 published HGSC scRNA-seq studies (**Table S1**) on 14,470 shared genes. Cells with < 500 UMIs, or <300 genes were removed, as were Scrublet-called doublets^63^, and artifact clusters. Samples with fewer than 500 remaining cells were removed. The atlas retained 1,980,703 cells from 362 samples of 158 patients. Data were integrated using scVI and scANVI^64^ (batch set per sample), seeded with CellAssign^65^ labels as scANVI annotations. Leiden clusters were annotated (**Figure S1-4, Table S2 – 4**).

#### Copy number

CopyKAT^66^ was run per sample, with a minimum of 200 epithelial cells and 50 reference cells. Fibroblast, T/NK, and endothelial cells were used as the diploid reference. Subclones (with 20 or more cells) were defined by hierarchical clustering and silhouette width.

#### Epithelial NMF and states

NMF (sckit-learn, k = 10) was fitted on a stratified set of 50,000 secretory cells and then applied to all 575,366 epithelial cells. The factor 2 (SecB) usage was used to classify secretory cells as SecA (<50^th^ percentile), Intermediate, or SecB (≥ 75^th^ percentile). In 250 samples with a minimum of 150 aneuploid epithelial cells, we recorded the principal component of malignant expression most correlated with SecB-SecA score as the dominant secretory axis of polarization.

#### Functional inference of epithelial states

PROGENy^67^ and DoRothEA^68^ activities (decoupler, multivariate linear model), Hallmark scores and scFEA^69^ flux (human M168 model) were averaged per epithelial state. Cell-cycle phase was assigned with scanpy. ConsensusOV^70^ subtypes were assigned to whole-sample pseudobulks of pre-treatment, primary samples (n = 96 samples, 77 patients). SecB vs SecA differential expression was tested with PyDESeq2 (design ∼sample + state, Wald test, BH) on sample-by-state pseudobulks (n = 261 samples, 135 patients).

#### Polarization scoring

Unless stated otherwise, polarization was the SecB – SecA UCell^71^ score on raw counts. SecA genes were MECOM, FBXO21, LGR5, LPAR3, PBX1, SOX17, and RCN2; SecB genes were KRT17, KRT19, KRT7, LCN2, PRSS22, SLPI and TACSTD2. They were selected for factor specificity, differential expression and cross-platform detectability. Each platform was scored in the gene space it shares with the atlas by UCell (Xenium, 457 genes; organoids, 14,271).TCGA-OV tumors were scored as the mean z-score of the 177 SecB -177 SecA NMF-specific genes. All gene sets are listed in **Table S6 and Table S9**. Visium HD bins were scored as the mean z-score of the SecB minus the SecA genes, and ICON7 arrays with UCell, each score z-scored across the 380 arrays. Organoid panels show the SecB UCell score.

#### Organoid snRNA-seq panel

PDOs were fixed and profiled with Parse Evercode WT v3. After doublet removal (scDblFinder) and QC (>500 UMIs, >300 genes, <20% mitochondrial reads), 10,455 cells remained.

#### Organoid experimental perturbations

OPTO98 organoids were cultured for 2,6,10 or 12 days of standard culture, with various environmental cytokines (IFNγ (20 ng/mL), TNFα (10 ng/mL), TGFβ (5 ng/mL), or WNT7A (500 ng/mL)), or eight varying medias (Basal media (BM, Advanced DMEM/F12, HEPES 10mM, antibiotic antimycotic 1X, B-27 supplement 1X, GlutaMax 1X, N-Acetyl-Cysteine 1mM), BM01 (BM + SB202190 0.5µM, A83-01 0.5µM, FGF-10 10ng/mL), BM02 (BM + FGF-4 10ng/mL), BM03 (BM + β-estradiol 100nM), BM1 (BM + Nicotinamide 5mM), BM2 (BM + EGF 5ng/mL, Heregulin-β-1 37.5ng/mL, hydrocortisone 0.5µg/mL, Forskolin 5µM), CGM (Advanced DMEM/F12, HEPES 10mM, antibiotic antimycotic 1X, B-27 supplement 1X, GlutaMax 1X, N-Acetyl-Cysteine 1.25mM, EGF 20ng/mL, β-FGF 100ng/mL, FGF-10 100ng/mL, Noggin 100ng/mL, Nicotinamide 1mM, Forskolin 10µM, β-estradiol 100nM, Y-27632 10µM), CGM Asc (CGM + cell free ascites fluid, 1:1) or HPLM (HPLM, HEPES 10mM, antibiotic antimycotic 1X, B-27 supplement 1X, GlutaMax 1X, N-Acetyl-Cysteine 1.25mM, EGF 20ng/mL, β-FGF 100ng/mL, FGF-10 100ng/mL, Noggin 100ng/mL, Nicotinamide 1mM, Forskolin 10µM, β-estradiol 100nM, Y-27632 10µM) for 6 days, with media replenished (in condition) every 3 days. At time of collection, organoids were subsequently dissociated using TrypLE for 45 min at 37°C and quenched with AdDMEM/F12. MULTI-seq barcoding, lipid-modified oligonucleotide anchor and barcode strands were prepared at a 1:1 molar ratio in serum-free PBS to generate sample-specific barcodes (2 µM, 10X anchor:barcode solution), alongside a 10X co-anchor solution (2 µM). Briefly, 50,000 cells per condition were washed twice and resuspended in 180 µL with PBS. Cells were incubated with 20 µL sample-specific 10X anchor: barcode solution on ice for 10 min with intermittent mixing every 3 min. This was followed by the addition of 22 µL of the 10X co-anchor solution and a further 10 min incubation with mixing occurring every 3 min on ice. Labeling was quenched with ice-cold 1% BSA-PBS, followed by two washes, filtration through a 40 µm strainer, and cell counting prior to downstream library preparation. These conditions were sample multiplexed in one 10X genomics library; after removing 4,017 doublets and 43 unassigned cells, 11,543 cells remained.

#### Organoid polarization quantification

We combined PDOs, atlas pseudobulk and three external datasets: DepMap organoids^72^, GSE299200^73^ and GSE284507^74^. All profiles were scored in one shared UCell space, with function defined as z(HIF) – z(proliferation). HIF: *NDRG1, SLC16A3, VEGFA, CA9, BNIP3, P4HA1, PDK1, SLC2A1;* Proliferation: *MKI67, TOP2A, PCNA, CCNB1, CDK1*.

#### KPCA.B tumor and Immunohistochemistry

One KPCA.B day 34 tumor was profiled by 10X Genomics Flex. After ambient-RNA removal and doublet filtering 20,598 cells remained, including 12,429 inferCNV-confirmed tumor cells. Tumor cells were scored with mouse orthologues of the polarization signature. De novo NMF programs were compared with human NMF factors by loading correlation with permutation null. KRT7 (clone: EPR17078) immunohistochemistry was performed on day 26 tumors. Briefly, FFPE sections were deparaffinized in xylene followed by tissue rehydration in decreasing levels of ethanol. Sections were rinsed twice in PBS and were subject to citrate-based heat induced antigen retrieval for 10 minutes followed by 10 minutes of tris-based retrieval. Slides were cooled, washed with PBS and treated with 3% hydrogen peroxide for 15 minutes. Slides were subsequently blocked with 2.5% horse serum for 30 minutes prior to overnight incubation with KRT7 (1:200). Slides were washed three times with PBS, incubated with rabbit secondary for 30 minutes. Slides were washed, incubated with 3,3-diaminobenzidine (DAB) for 90 seconds and counterstained with hematoxylin. Slides were dehydrated with increasing levels of ethanol, mounted with coverslips and scanned (Zeiss Z1; 20x, 0.22µm per pixel); images were exported using QuPath.

#### Xenium

We designed a custom 477-gene Xenium panel (v1 chemistry, **Table S7**). Probes were benchmarked against the atlas and 28 discordant genes were excluded from use in annotation. Cells were retained with 10 or more transcripts, 5 or more genes and an area of 10 - 500µm^2^ and <5% control-probe counts. TMA cores were de-arrayed, QC filtered and pooled per patient, and patients with 500 or more cells were retained for spatial and survival analyses. The final data comprise 1.9 million cells from 8 whole-tissue sections, 1.5 million TMA core cells from 503 cores, representing 378 patients and 34,281 fallopian tube epithelium cells from 15 patients. Cells were annotated automatically using SingleR using the atlas as the reference dataset. Secretory cells were split into SecA, Intermediate and SecB, resulting in 18 final cell types. Pathway modules were scored throughout using UCell.

#### Spatial statistics

SecA-SecB spatial association was quantified by global bivariate Moran’s I, between each secretory cell’s SecA score and the SecB scores of secretory cells within 50 µm. Weights were binary with 999 permutations per tissue and 199 per TMA core (minimum of 500 secretory cells). Per cell, it was quantified by bivariate LISA (499 permutations, BH within tissue). From cells of interest, we derived local measures of density, neighborhood pathway scores, local epithelial hypoxia or polarization scores and distance to the nearest pericyte or endothelial cell. Gradients were fitted with tissue-weighted GAMs (mgcv, s(x, k = 10) over all cells of the eight whole tissues, and immune composition with per-tissue binomial GAMs. Curves span the 1st–99th percentiles. Each panel was tested per tissue by comparing the top and bottom 20% (paired Wilcoxon, n = 8). Nuclear and cell areas are derived from Xenium segmentation. Macrophage morphology was determined in tissues or TMA patients with a minimum of 10 macrophages per tertile of the lowest and highest niche polarization. Local cell density was the number of cells within 50 µm of each secretory cell per mm² of the 50-µm circle (median per sample and state).

#### Visium HD

WT01 was profiled by Visium HD and analyzed in 16 and 8 µm bins. Bins were classified as epithelial or macrophage by marker UMI counts, and bin polarization was the mean z-score of the SecB minus the SecA genes. Macrophage bins with 2 or more SecB high epithelial bins within 48 µm were compared to distal macrophage bins.

Fold changes were adjusted for macrophage content and genes attributed to adjacent epithelium were excluded.

#### Macrophage niches

In Xenium, SecB-niche macrophages had 5 or more SecB cells within 50 µm, controls had no SecB cells and a minimum of 5 SecA cells. They were compared per tissue. SecB cells within 30µm of niche macrophages were compared to SecB cells with no surrounding macrophages. For each macrophage with ≥3 secretory neighbors within 50 µm, the SecB share of those neighbors was compared between the top (hypoxic–glycolytic) and bottom (tissue-resident) tertiles of a program score. The score was nine hypoxia–glycolysis genes minus four resident genes (**Table S9**). The comparison was made per whole tissue and per TMA patient (n = 311).

#### Protein correlation

Previously published immunofluorescence intensities from TMA A were correlated per core (Pearson and Spearman; 177 final TMA A cores from 95 patients, 157–163 with IF for each marker) with Xenium transcript expression and mean polarization.

#### Survival

TMA survival analyses included patients with primary, pre-treatment HGSC and represented by two tumor cores. Patients given neoadjuvant chemotherapy (n = 5) or with non-HGSC histology on central review (n = 4) were excluded. Follow-up was censored at 5 years, and stage, residual disease and age had to be complete (OS n = 104, 40 events; PFS n = 103, 71 events). Kaplan–Meier curves compared patients in the lowest and highest tertiles of per-patient polarization, the middle tertile left out (log-rank test). SecA proportion, SecB proportion and mean polarization were z-scored and each tested in a Cox model adjusted for stage, residual disease and age, with Benjamini–Hochberg FDR across these six tests (**Table S11**). Proportional hazards were assessed with Schoenfeld residuals (**Table S11**). TCGA-OV polarization was compared between the highest and lowest tertiles by log-rank test and tested in Cox models adjusted for epithelial fraction (BayesPrism), stage and age. In ICON7 (GSE140082, stage III/IV, n = 191), tumors were split at the median polarization. The bevacizumab effect on PFS was estimated per group, with a 15-month time split. The treatment x polarization interaction was tested by likelihood-ratio test in a Cox model adjusted for adjusted for FIGO stage, debulking status and age. In each pair of forest plots, q is the Benjamini–Hochberg FDR across the estimates shown.

### STATISTICAL ANALYSES

Inference used independent units: patients or samples (atlas level conclusions), models (PDOs), sections or TMA patients (Xenium) and patients (survival). Cell- and bin-level analyses are descriptive. Paired comparisons used two-sided Wilcoxon signed-rank tests. Other tests were Kruskal–Wallis tests, DESeq2 Wald and limma-voom moderated t-tests on pseudobulks, permutation tests, Pearson and Spearman correlations, log-rank tests and Cox regression; all tests were two-sided. Benjamini–Hochberg FDR was applied within stated families (genes in each differential-expression analysis, BiLISA cells within each tissue, the KPCA.B loading correlations **of Figure S10**, and the Cox model sets of **Figure 7** and **Tables S11–S12**); other p values are nominal. Specific test, sample n value and error definitions are defined within figure legends. Analyses used R 4.4.3 (Bioconductor 3.20) and Python 3.12.9; package versions are specified within the repository.

## Declaration of generative AI and AI-assisted technologist in the manuscript preparation process

During the preparation of this work, the authors used Claude Opus 4.8-5.5 for coding and manuscript review. The author(s) reviewed and edited the output as needed and take full responsibility for the content of the published article.

## SUPPLEMENTARY TABLES

**Table S1. Characteristics of the 13 scRNA-seq studies integrated in the HGSC atlas.** Per study publication, data accession, sequencing platform and chemistry, tissue processing, and the number of patients, samples, sequencing libraries, and cells retained in the final atlas (1,980,703 cells, 158 patients, 362 samples), with anatomic site and treatment status composition. Additional sheets list the metadata for each sample including clinical annotations where available (treatment status, stage, anatomic site, metastatic site, age, treatment response, BRCA status, HRD status, TP53 status) and cell counts and the number of cells per study for each level-1 lineage.

**Table S2. Differentially expressed markers for cell type level 1 annotation.** Wilcoxon rank-sum test results for upregulated genes in each of the 13 level 1 cell type compartments versus all other cells (p adj < 0.05, log2FC > 0). Columns indicate cell type, rank, gene, Wilcoxon score, log2 fold change, p-value and adjusted p-value (Benjamini-Hochberg). 51,592 significant gene-cell type pairs across annotated cells; the top 50 markers per cell type are listed in a separate sheet.

**Table S3. Differentially expressed markers for cell type level 2 annotation.** Wilcoxon rank-sum test results for upregulated genes in each level 2 cell type versus the other level 2 clusters of the same level 1 compartment (p adj < 0.05, log2FC > 0). Columns indicate level 1 compartment, level 2 cell type, rank, gene, Wilcoxon score, log2 fold change, p-value and adjusted p-value (Benjamini-Hochberg). 197,160 significant gene-cell type pairs across 49 level 2 cell types; pericytes and neutrophils comprise a single level 2 type and are represented in Supplementary Table 2. The top 50 markers per level 2 cell type are listed in a separate sheet.

**Table S4. Differentially expressed markers for ciliated and secretory clusters.** Ciliated and secretory epithelial cell clusters were compared by Wilcoxon rank-sum test. Columns indicate gene, log2 fold change, Wilcoxon score, p-value, adjusted p-value (Benjamini-Hochberg), direction of enrichment and percent expression within each group. 10,029 significant DEGs (p adj < 0.05).

**Table S5. Non-negative matrix factorization (NMF) programs of the atlas epithelial compartment.** NMF (k = 10) was performed on 3,007 genes (3,000 highly variable genes plus signature genes). Sheets give the annotation of each factor and its top genes; the loading of every gene in every factor, with its within-factor rank and specificity; the top 50 genes per factor; the classification of genes as SecA-specific (177), SecB-specific (177) or shared (124), by the 90th percentile of the Factor 3 and Factor 2 loadings; and consensus metaprograms from per-patient consensus NMF (k = 5–10, 30 runs per k). Factor 3 corresponds to the SecA program and Factor 2 to the SecB program.

**Table S6. SecA and SecB gene signatures used for polarization scoring.** The seven SecA genes (MECOM, FBXO21, LGR5, LPAR3, PBX1, SOX17, RCN2) and seven SecB genes (KRT17, KRT19, KRT7, LCN2, PRSS22, SLPI, TACSTD2) are listed with their NMF loadings and ranks, SecB versus SecA pseudobulk DESeq2 log2 fold change and adjusted p (**Figure 2g**), the percentage of atlas SecA, Intermediate and SecB cells expressing each gene, Xenium panel membership and probe quality, and the analyses in which each signature was used. Additional sheets list the scoring method for each dataset and the 177 SecA-specific and 177 SecB-specific genes used to score TCGA-OV.

**Table S7. Xenium gene panel composition and cross-platform probe quality control.** The custom Xenium panel of 477 genes is listed with its design rationale, cell type marker assignment, SecA/SecB signature and pathway module membership, and probe QC metrics against the scRNA-seq atlas: correlation of gene–gene co-expression profiles, top-20-partner Jaccard overlap, mean and variance ratios, detection-rate and non-zero-mean differences, and overdispersion. Genes with ≥3 QC flags failed (31 genes). Of these, 28 were excluded from SingleR annotation (429 genes used for annotation), and three essential markers were retained with flags. All genes were retained for expression and pathway analyses; 20 genes absent from the atlas are marked Xenium-only.

**Table S8. Clinical and sample characteristics of the Xenium spatial transcriptomics cohort.** De-identified characteristics are given for the 8 whole HGSC tissue sections (treatment status: 4 treatment-naïve, 4 post-chemotherapy; FIGO stage; histology; cells profiled; epithelial composition), for the patients on the two HGSC tissue microarrays (378 patients with ≥500 cells across 503 cores), and for the 15 fallopian tube epithelium cores. TMA patient data include age band, FIGO stage, residual disease, neoadjuvant treatment, histology, grade, cores and cells profiled, OS and PFS (months) with event status, and inclusion in the Figure 7 survival analyses (OS n = 104, PFS n = 103). Study codes do not correspond to any clinical or biobank identifier.

**Table S9. Gene sets used for pathway scoring of the Xenium spatial data.** The 37 custom UCell pathway modules (325 gene entries) are listed with their category, genes and probe QC status. Also listed are the custom epithelial (DNA damage repair, epigenetic silencing; Figure 5a) and macrophage (glycolytic, NF-κB activated, tissue-resident identity, chemokine; Figure 5i) gene sets, and the two gene sets of the Figure 6g macrophage program score (glycolytic and tissue-resident; scored as z-scored expression, not UCell), together with the scoring method for each set and the figures in which it is reported.

**Table S10. Differential expression of macrophages in SecB niches.** Visium HD macrophage bins (WT01, 16 µm and 8 µm) adjacent to SecB-high epithelium (≥2 SecB-high bins within 48 µm) were compared with distal macrophage bins (Figure 6e). The log2 fold change of mean CP10k is given before and after adjustment for macrophage content (median log2 fold change of pan-macrophage genes subtracted). Each gene is classified as macrophage-attributable, SecB epithelial carry-over or stromal, and bin-level Mann– Whitney p values with Benjamini–Hochberg FDR are reported descriptively. Xenium SecB-niche macrophages (≥5 SecB epithelial cells within 50 µm) were compared with control macrophages (no SecB and ≥5 SecA cells within 50 µm) by pseudobulk limma-voom paired by section, across 7 whole tissues with ≥50 macrophages per group (WT07 had 39 SecB-macrophages and was excluded). Per-section log2 fold changes and consistency are included.

**Table S11. Multivariable Cox regression of secretory polarization in the Xenium TMA cohort.** For patients of the final TMA cohort represented by two final tumor cores (OS n = 104, 40 events; PFS n = 103, 71 events), each of the three secretory measures (SecA and SecB proportion of secretory cells; mean polarization score) was modeled separately, adjusted for FIGO stage (III/IV vs I/II), residual disease (suboptimal vs optimal) and age, for 5-year OS and PFS (Figure 7c,d). Hazard ratios are per 1 SD increase, with 95% confidence intervals, Wald p, Benjamini–Hochberg FDR across the six tests of Figure 7c,d, all covariate estimates and Schoenfeld-residual proportional-hazards tests. The Kaplan–Meier comparison of the lowest and highest polarization tertiles in Figure 7a,b (tertile cut-offs, group sizes, events, 5-year survival and log-rank p) is included.

**Table S12. Cox regression of secretory polarization and bevacizumab benefit in the ICON7 trial.** Pre-treatment tumors from FIGO stage III/IV patients (GSE140082; n = 191) were classified as SecA-polarized (n = 96) or SecB-polarized (n = 95) at the cohort median polarization score. Every model is adjusted for FIGO stage, debulking status and age. The table reports the association of continuous polarization (per 1 SD) with OS and PFS within each treatment arm; the effect of bevacizumab on PFS within each polarization group (Figure 7k,l); the time-varying bevacizumab effect before and after 15 months, with the period interaction (Figure 7m,n); and the treatment x polarization-group interaction for PFS, with its likelihood-ratio test. Kaplan–Meier median survival, group sizes and all model coefficients are included. Benjamini–Hochberg q values for Figure 7m,n and for the period interaction are included.

